# The Amyloid Precursor-like Protein APL-1 Regulates Axon Regeneration

**DOI:** 10.1101/305284

**Authors:** Lewie Zeng, Rachid El Bejjani, Marc Hammarlund

**Affiliations:** Department of Genetics, Yale University School of Medicine, New Haven, CT 06520.; Biology Department, Davidson College, Davidson, NC 28035; Department of Neuroscience, Yale University School of Medicine, New Haven, CT 06520.

## Abstract

Members of the Amyloid Precursor Protein (APP) family have important functions during neuronal development. However, their physiological functions in the mature nervous system are not fully understood. Here we use the *C. elegans* GABAergic motor neurons to study the post-developmental function of the APP-like protein APL-1 in vivo. We find that *apl-1* has minimum roles in the maintenance of gross neuron morphology and function. However, we show that *apl-1* is an inhibitor of axon regeneration, acting on mature neurons to limit regrowth in response to injury. The small GTPase Rab6/RAB-6.2 also inhibits regeneration, and does so in part by maintaining protein levels of APL-1. To inhibit regeneration, APL-1 functions via the E2 domain of its ectodomain; the cytoplasmic tail, transmembrane anchoring, and the E1 domain are not required for this function. Our data defines a novel role for APL-1 in modulating the neuronal response to injury.

## Introduction

The Amyloid Precursor Protein (APP) belongs to a family of type I, single pass transmembrane proteins that are highly conserved from mammals to *C. elegans*, and that have prominent expression in the nervous system (Hornsten et al., 2007; Lorent et al., 1995; Luo et al., 1990; Wiese et al., 2010). Some of the studied family members include APP, APLP1, and APLP2 in mammals (Goldgaber et al., 1987; Kang et al., 1987; Slunt et al., 1994; Tanzi et al., 1987; Wasco et al., 1992, 1993); APPa, APPb, APLP1, and APLP2 in zebrafish (Liao et al., 2012; Musa et al., 2001); APPL in *Drosophila* (Luo et al., 1990; Rosen et al., 1989); and APL-1 in *C. elegans* (Daigle and Li, 1993). Although these proteins are best known for the role of human APP in Alzheimer’s disease, the Aβ peptide sequence implicated in the pathogenesis of Alzheimer’s is only present in APP itself, and is not found in mammalian APLP1and APLP2, or in *C. elegans* or *Drosophila*, suggesting it is a more recent addition in evolution. The neuronal expression and the overall conservation of the APP family outside the Aβ sequence suggests that these genes have important physiological functions in neurons beyond the pathogenic role of Aβ.

Genetic models have been used to analyze the functions of the APP family in neurons. Loss of function studies in mice demonstrated that the APP family is essential and partially redundant: while the single knockouts and *App/Aplp1* double knockouts are viable, *App/Aplp2* and *Aplp1/Aplp2* double knockouts and the triple knockouts die within a few days after birth (Heber et al., 2000; von Koch et al., 1997; Zheng et al., 1995). The single ortholog of *App* in *C. elegans, apl-1*, is also an essential gene (Hornsten et al., 2007; Wiese et al., 2010). By contrast, *Appl* mutants in *Drosophila* are viable (Luo et al., 1992), and the effect of multiple knockouts in zebrafish is not known. Despite these differences, however, analysis of the nervous system of mutant animals has revealed a conserved function for the APP family in multiple aspects of neuronal development. Defects in synaptogenesis, axon outgrowth, and cell migration have been described across mouse, fly, and fish models (Dawson et al., 1999; Herms et al., 2004; Luo et al., 1992; Phinney et al., 1999; Rama et al., 2012; Seabrook et al., 1999; Soldano et al., 2013; Song and Pimplikar, 2012; Torroja et al., 1999; Wang et al., 2005, 2009; Yang et al., 2005; Young-Pearse et al., 2007). In addition, their effect on axon outgrowth has also been studied in cultured cells and explants (Allinquant et al., 1995; Araki et al., 1991; Jin et al., 1994; Milward et al., 1992; Ninomiya et al., 1994; Ohsawa et al., 1995; Perez et al., 1997; Rama et al., 2012; Wallace et al., 1997; Young-Pearse et al., 2008). Thus, the APP family is critical for normal development. However, although APP family members are indeed expressed during development (Clarris et al., 1995; Löffler and Huber, 1992; Moya et al., 1994), they are also expressed across species at high levels in the mature nervous system (reviewed in Müller et al., 2017). Little is known about how these critical neuronal proteins function in the mature nervous system.

Here, we analyze the post-developmental function of *apl-1* in the *C. elegans* nervous system. We find that although *apl-1* is not required for gross nervous system structure or function under normal conditions, it acts following axon injury to inhibit axon regeneration. Levels of the APL-1 protein and inhibition of regeneration are dependent on retrograde recycling of the protein by the small GTPase RAB-6.2. Further, we show that inhibition of regeneration by APL-1 uses an extracellular mechanism, acting independently of APL-1’s cytoplasmic sequence and its transmembrane domain. Our findings identify a novel function for APL-1 as an inhibitor of axon regeneration, operating outside injured neurons to affect their ability to regenerate.

## Results

### APL-1 Inhibits Axon Regeneration in the GABAergic Motor Neurons

To assess the physiological role of APP family proteins in mature neurons, we used the GABAergic motor neurons in *C. elegans* as a model (Figure 1A). The single *C. elegans* APP ortholog, *apl-1*, is known to be expressed in various cell types including neurons, hypodermis, and a number of junction cells (Hornsten et al., 2007; Wiese et al., 2010). We examined *apl-1* expression using a *Papl-1*∷mCherry reporter driven by 6.3 kb of the *apl-1* promoter, in the *juIs76* strain expressing GFP under the control of a GABAergic neuron-specific promoter *Punc-25* (Jin et al., 1999). We observed *Papl-1*∷mCherry in all GABAergic motor neurons (Figure 1B). Expression was also observed in other ventral nerve cord neurons, head and tail neurons, sublateral nerve bundles, uterine cells, and the hypodermis, consistent with previous reports (Hornsten et al., 2007; Wiese et al., 2010). We also examined the expression of an APL-1-GFP protein fusion driven by the same *Papl-1* promoter sequence in a strain expressing mCherry under the *Punc-47* (McIntire et al., 1997) GABAergic neuron-specific promoter. This fusion construct was expressed in the same cells as *Papl-1*∷mCherry, and fluorescence was localized to subcellular compartments that include the ER and other vesicular compartments (Figure 1C, S1B), consistent with the subcellular localization of mammalian APP family members which are found in the ER, endosomes, Golgi, and plasma membrane (Kaden et al., 2009). It has been previously reported that C-terminally tagged APL-1-GFP expressed under a *Papl-1* promoter is capable of rescuing the phenotype of *apl-1(yn10* and *tm385)* mutants (Hornsten et al., 2007; Wiese et al., 2010).

**Figure 1.**
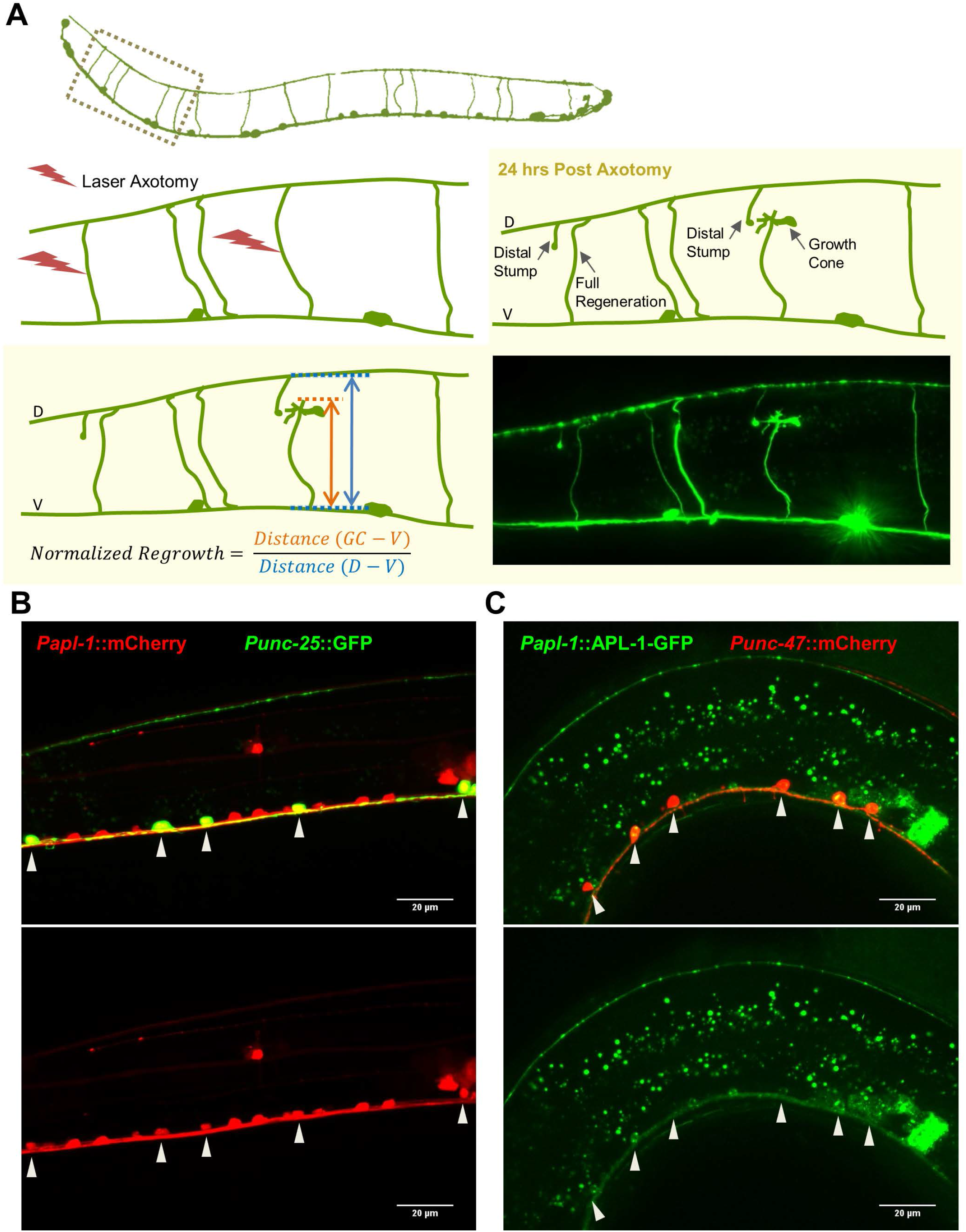
APL-1 is expressed in the GABAergic motor neurons. (A) Diagram of GABAergic motor neurons in *C. elegans* and the method for scoring axon regeneration. GABAergic axon commissures in L4 stage worms are severed in the middle using a 435nm pulsed laser. 24 hours (at 20°C) post axotomy, regeneration of each individual axon is scored as “normalized regrowth”, which is calculated as the distance between the front edge of the growth cone and the ventral nerve cord divided by the distance between the dorsal and ventral nerve cords at the same position. (B) Transcriptional reporter of *apl-1* (6.3kb promoter region) driving mCherry. Expression seen in GfP labeled *(unc-25* promoter) GABAergic motor neurons *(juIs76* background) (arrowheads). Scale bar 20μm. (C) Translational reporter of *apl-1.* APL-1 protein tagged with GFP under 6.3kb *apl-1* promoter. Expression seen in mCherry labeled *(unc-47* promoter) GABAergic motor neurons *(wpls40* background) (arrowheads). Scale bar 20μm. (Z-projection of a thin section of the worm was taken to show the ventral nerve cord clearly; the dorsal nerve cord is not shown in this picture.)

A major challenge to studying the post-developmental functions of *apl-1* in *C. elegans* is that *apl-1* is an essential gene required during larval development. *apl-1* null worms arrest at L1/L2 early larval stages, possibly due to molting defects (Hornsten et al., 2007; Wiese et al., 2010). In order to investigate the function of *apl-1* in mature GABAergic motor neurons, we built a conditional allele based on the Cre-*loxP* system. We used CRISPR/Cas9 mediated gene editing (method adapted from Arribere et al., 2014; Dickinson et al., 2013; Farboud and Meyer, 2015; Friedland et al., 2013; Paix et al., 2014) to insert *loxP* sites flanking the endogenous *apl-1* coding sequence (Figure 2A). We refer to this allele as *apl-1(floxed).* We expressed GABAergic motor neuron-specific Cre recombinase driven by the *Punc-47* promoter in *apl-1(floxed)* animals, and validated, using single worm PCR, that the cell type-specific Cre recombinase expression is able to excise the entire *apl-1* gene coding region between the two *loxP* sites (Figure 2A). Thus, these animals, which we refer to as *Pgaba*∷Cre; *apl-1(floxed)*, are a new model for the *in vivo* characterization of the function of APP family proteins in mature neurons.

**Figure 2.**
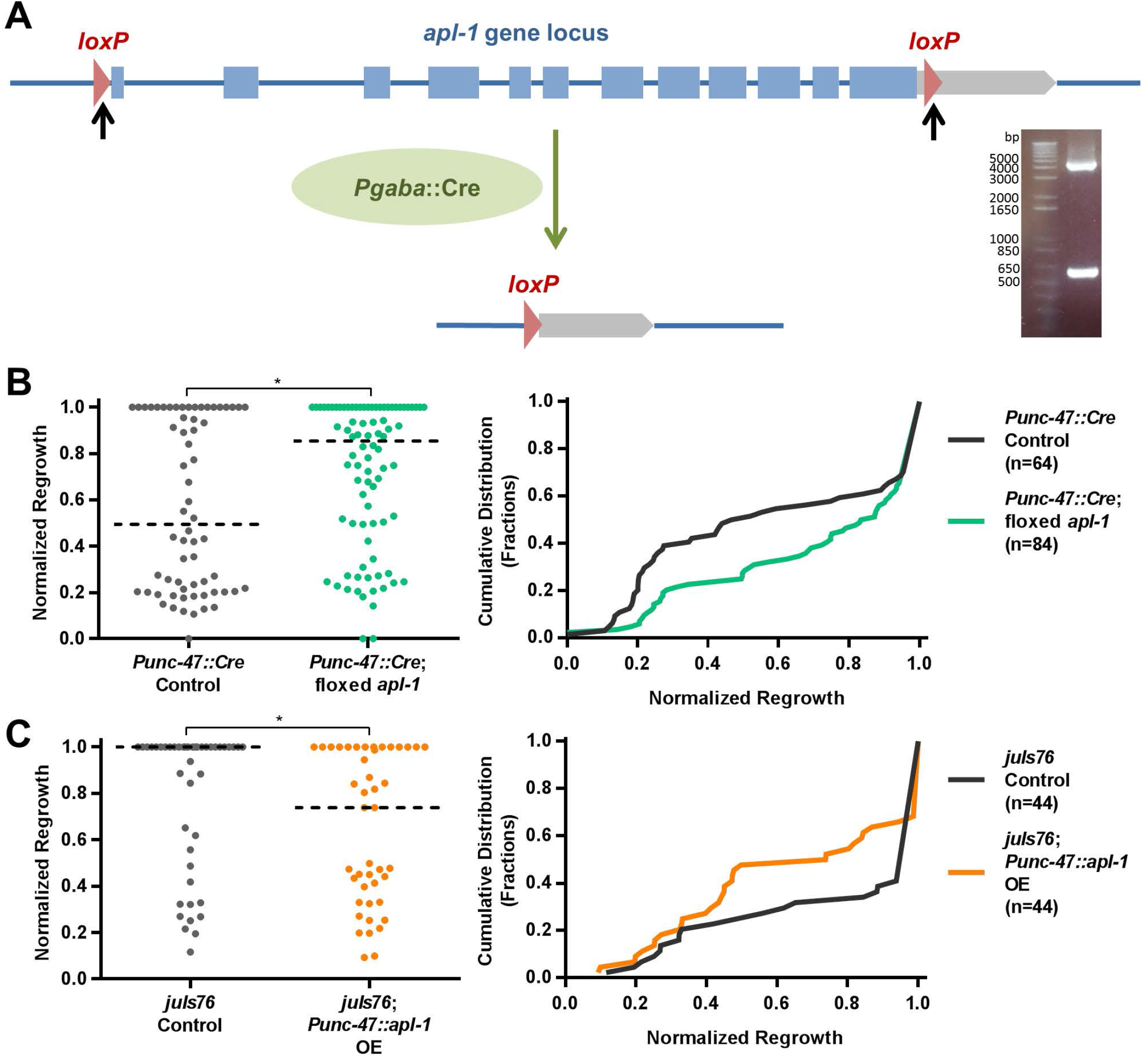
*apl-1* inhibits axon regeneration in the GABAergic motor neurons. (A) Diagram showing sites of *loxP* insertions (black arrows) at the endogenous *apl-1* gene locus, and excision by Cre recombinase. Gel shows an example of single worm PCR across the genomic locus. The bigger band corresponds to the unexcised locus, and the smaller band corresponds to the locus after excision in cells that express Cre. (B) Axon regeneration is increased in the GABA specific Cre-loxP mediated *apl-1* conditional knockout (cKO) worm compared to the Cre only control background *(wpIs78*, Cre under *unc-47* promoter). Left panel is a scatter plot of the normalized regrowth of injured axons, each dot representing the value for one axon. Dashed line indicates the median. Right panel plots the cumulative distribution curves of the data in the left panel. Right shifting of the curve indicates increased regeneration. The nonparametric Kolmogorov-Smirnov test is used to compare the cumulative distributions. (Kolmogorov-Smirnov D=0.2619; P=0.0137; the number of axons scored “n” is shown on graph.) The same type of plots and statistic test are used for all following regeneration data. (C) Axon regeneration is decreased in worms overexpressing APL-1 specifically in the GABAergic motor neurons *(unc-47* promoter) compared to *juIs76* control marker background. (Kolmogorov-Smirnov D=0.2955; P=0.0429)

APP family members are expressed in neurons throughout life; indeed, the continued expression of APP is likely a precondition for its role in Alzheimer’s disease. This suggests that APP proteins have functions in mature neurons, after development has ended. To discover such potential functions, we compared the morphology and function of GABAergic motor neurons in *Pgaba*∷Cre; *apl-1(floxed)* animals to controls. We found that expression of Cre alone caused a low level of developmental axon guidance or outgrowth defects in *Pgaba*∷Cre control worms. These defects were not increased in *Pgaba*∷Cre; *apl-1(floxed)* worms (Figure S2B). Because Cre expression under control of the GABAergic promoter only slightly precedes axon outgrowth, these experiments do not rule out a potential role for APL-1 in GABAergic motor neuron development. The lack of developmental defects provides a clean background for the study of post-developmental roles for APL-1.

To assess the function of mature neurons that lack APL-1, we examined young 1-day adult worms’ resistance to hypercontracted paralysis induced by an acetylcholinesterase inhibitor aldicarb, which can provide a sensitive measure of GABAergic motor neuron synaptic function: animals that release less GABA are hypersensitive to aldicarb, presumably because the inhibitory effects of GABA normally offset some of the excitatory effects of aldicarb (Miller et al., 1996; Vashlishan et al., 2008). We found that *Pgaba*∷Cre; *apl-1(floxed)* worms were slightly more resistant to aldicarb-induced paralysis compared to *Pgaba*∷Cre control worms, suggesting increased GABA release (Figure S2E). These data suggest that in GABA neurons, APL-1 has a small inhibitory effect on vesicle release. We also examined the effect of aldicarb application on two viable *apl-1* domain-deletion mutants, *apl-1*Δ*TM-ICD* and *apl-1*Δ*E1*, which we generated by CRISPR/Cas9 mediated gene editing (discussed later). *apl-1*Δ*TM-ICD* mutant worms exhibited hypersensitivity to aldicarb-induced paralysis, whereas *apl-1*Δ*E1* mutants were similar to wildtype controls (Figure S2F). These results, which affect all *apl-1* rather than just *apl-1* in GABA neurons, are consistent with previously reported results for *apl-1* RNAi worms, which are hypersensitive to aldicarb (Wiese et al., 2010). Together, the analysis of aldicarb resistance suggests that GABA neuron-specific loss of *apl-1* has only minor effects on function, but leaves open the possibility that *apl-1* has greater functional roles in other neuron types.

Next, we considered the possibility that *apl-1* functions in aging neurons to maintain normal morphology. GABAergic motor neurons and other neurons in *C. elegans* accumulate low levels of morphological defects with advancing age, such as bead-like aggregations and small branch-like protrusions (Pan et al., 2011; Tank et al., 2011; Toth et al., 2012). We examined age-related axon morphology phenotypes in the *Pgaba*∷Cre; *apl-1(floxed)* worms compared to *Pgaba*∷Cre control worms. There was no difference in the fraction of axons forming bead-like aggregations or small branch-like protrusions (Figure S2C, S2D). In general, we observed no morphological effect of loss of *apl-1* in aging animals, similar to the lack of effect in young adult animals (see above). We conclude that loss of *apl-1* has at most minimal effects on basal neuronal structure and morphology, even in aging animals.

Finally, we examined the effect of loss of *apl-1* on axon regeneration. Axon regeneration represents a dramatic instance of neuronal plasticity, in which neurons must respond to axon injury by re-entering a growth-competent state, and then must successfully drive growth cones through a mature environment toward distant targets. Unexpectedly, we found that axon regeneration of the GABAergic motor neurons was improved in the *Pgaba*∷Cre; *apl-1(floxed)* animals compared to the *Pgaba*∷Cre control (Figure 2B, S2A). These data point to a new function for APL-1, in inhibiting axon regeneration. To further test this idea, we overexpressed the full-length APL-1 in the GABAergic motor neurons under the promoter *Punc-47*, and we found that axon regeneration was decreased in these neurons (Figure 2C). Together, the conditional loss of function and the overexpression data suggest that the neuronal APL-1 inhibits axon regenerative capacity in the GABAergic motor neurons.

### Rab-6.2 Maintains Regeneration-Inhibitory Levels of APL-1

To determine how expression of the APL-1 protein might contribute to reduced regeneration, we expressed APL-1-GFP (Figure 3A) under the *Punc-47* promoter, and examined its localization in the GABAergic motor neurons. In uninjured neurons, APL-1-GFP is localized in subcellular compartments in the cell body and along the axons (Figure 3A, S1A), similar to the subcellular localization pattern for mammalian APP in neurons (Caporaso et al., 1994; Ferreira et al., 1993; Moya et al., 1994; Simons et al., 1995; Tienari et al., 1996). Live imaging demonstrated that APL-1-GFP is trafficked within axons (Figure S1A), again similar to its behavior in mammalian neurons (Koo et al., 1990; Sisodia et al., 1993). We also looked at APL-1-GFP localization 8 to 10 hours after laser axotomy. For non-regenerating axons, APL-1-GFP can be seen accumulating at injured axon tips (Figure S3A), possibly due to interrupted axonal transport at the injured site. Similar accumulation of APP has been observed in injured mammalian neurons (Gentleman et al., 1993; Otsuka et al., 1991; Pierce et al., 1996); in fact, APP has often been used as a marker for axonal injury (summarized in Stone et al., 2000). For the remaining distal axon segment, APL-1-GFP can be seen accumulating only at the swollen severed tip and absent from the more distal portion (Figure S3A), possibly reflecting the retrograde axonal transport and degradation of APL-1-GFP and the absence of arrival of newly synthesized APL-1-GFP from the cell body. For regenerating axons, APL-1-GFP is present all along the axon and also in the growth cone area (Figure S3A), consistent with the localization of APP in cultured mammalian neurons (Sabo et al., 2003; Yamazaki et al., 1995).

**Figure 3.**
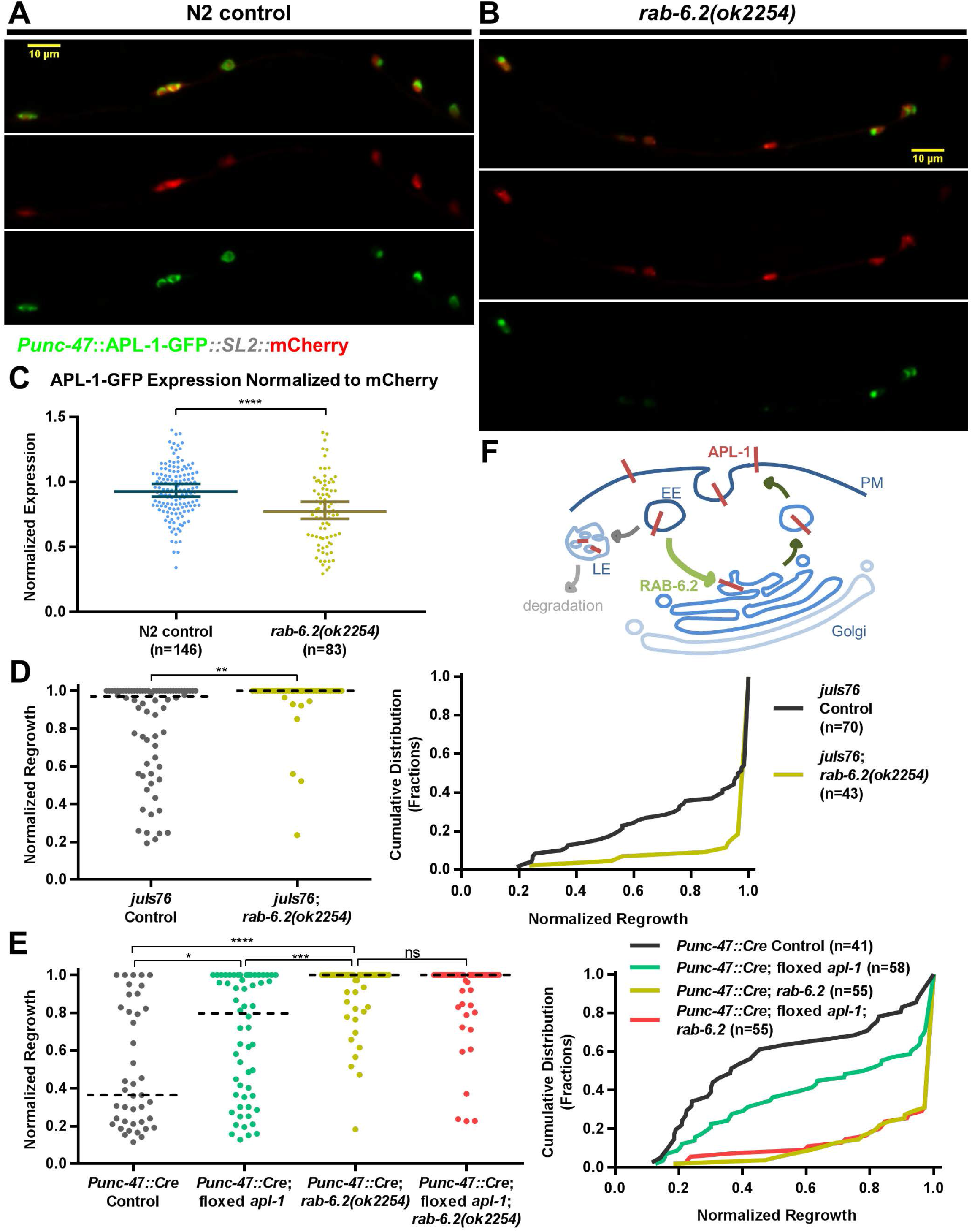
APL-1 depends on *rab-6.2* to maintain its protein level and inhibit axon regeneration in GABAergic motor neurons. (A) APL-1-GFP expressed in GABAergic motor neurons *(unc-47* promoter) of N2 control worms, along with mCherry separated by *SL2* splice leader sequence *(gpd-2/gpd-3* intergenic region) for trans-splicing and co-expression. Scale bar 10μm. (B) Same expression array as (A) in *rab-6.2(ok2254)* null mutant background. Scale bar 10μm. (C) Quantification of APL-1-GFP expression in cell bodies normalized to mCherry comparing N2 control (A) and *rab-6.2(ok2254)* null (B) backgrounds. Mann-Whitney U test is used (control median=0.9277; rab-6.2 median=0.7717; P<0.0001). Line and error bar shows median and 95% CI. APL-1-GFP level is lower in *rab-6.2(ok2264)* mutant. (D) Axon regeneration is increased in *rab-6.2(ok2254)* mutant compared to *juIs76* control. (Kolmogorov-Smirnov D=0.3568; P=0.0023) (E) Axon regeneration is increased more in *rab-6.2(ok2254)* mutant than in apl-1 Cre-loxP GABA cKO compared to Cre only control *(wpIs78)*, while double mutant of *rab-6.2(ok2254)* and *apl-1* cKO does not further increase regeneration than *rab-6.2(ok2254)* single mutant. (control vs *apl-1* cKO Kolmogorov-Smirnov D=0.2822, P=0.0436; control vs *rab-6.2* Kolmogorov-Smirnov D=0.5929, P<0.0001; *apl-1* cKO vs *rab-6.2* Kolmogorov-Smirnov D=0.3987, P=0.0003; *rab-6.2* vs *rab-6.2* and *apl-1* cKO double Kolmogorov-Smirnov D=0.05455, P>0.9999) (F) Model of RAB-6.2 function in promoting retrograde recycling from the early endosomes to the trans-Golgi network. (PM, plasmamembrane; EE, early endosomes; LE, late endosomes)

Rab6 is known to affect the trafficking of APP in mammalian cells (McConlogue et al., 1996; Thyrock et al., 2013). A mammalian Rab6 isoform has been shown to mediate protein transport from early and recycling endosomes to the trans-Golgi network (TGN) (Mallard et al., 2002; Young et al., 2005). In *C. elegans* neurons, the Rab6 worm ortholog RAB-6.2 has been shown to mediate the retrograde trafficking and recycling of transmembrane proteins GLR-1 and MIG-14 from the early endosomes back to the TGN (Zhang et al., 2012, 2016). We hypothesized that in *C. elegans* GABAergic motor neurons, RAB-6.2 acts after endocytosis to promote the recycling of APL-1 via retrograde trafficking, diverting it from the protein degradation pathway (Figure 3F). To test whether RAB-6.2 affects APL-1 protein levels, we examined the expression of APL-1-GFP in the GABAergic motor neurons of wildtype and *rab-6.2(ok2254)* null mutant worms. We found that at L4 stage or older, the abundance of APL-1-GFP was significantly decreased in the *rab-6.2(ok2254)* null mutant GABA motor neurons (Figure 3A, 3B, quantification in 3C). These data indicate that RAB-6.2 is required to maintain APL-1 protein levels in the GABAergic motor neurons. To determine whether RAB-6.2 acts after endocytosis of APL-1, we expressed APL-1ΔC-GFP, deleting the cytoplasmic C-terminus of APL-1, which contains the conserved motifs required for APP endocytosis (Koo and Squazzo, 1994; Perez et al., 1999). We found that APL-1ΔC-GFP levels were no longer decreased by the loss of RAB-6.2, consistent with the model that RAB-6.2 acts after APL-1 endocytosis (Figure S4A, S4B, S4C). To confirm and extend these results, we inserted a 3xFlag tag at the N-terminus of the endogenous APL-1 just after the signal peptide domain via CRISPR/Cas9 mediated gene editing of the *apl-1* gene locus (Figure 4A). The resulting fusion protein reports on APL-1 levels everywhere it is expressed, rather than specifically in the GABAergic motor neurons. We confirmed by western blot that the anti-Flag antibody detected a band at over 100kD in the 3xFlag tagged worms, representing the full-length 3xFlag-APL-1, whereas the band was not present in the untagged negative control worms (Figure S4D). Next, we examined 3xFlag-APL-1 protein expression in *rab-6.2* null mutants, and found that it was significantly decreased. These data confirm that RAB-6.2 is required to maintain APL-1 protein levels. In addition, because the GABAergic motor neurons represent only a fraction of overall APL-1 expression (Figure 1C), these data suggest that the effect of RAB-6.2 on APL-1 abundance extends to other cells in addition to the GABAergic motor neurons.

**Figure 4.**
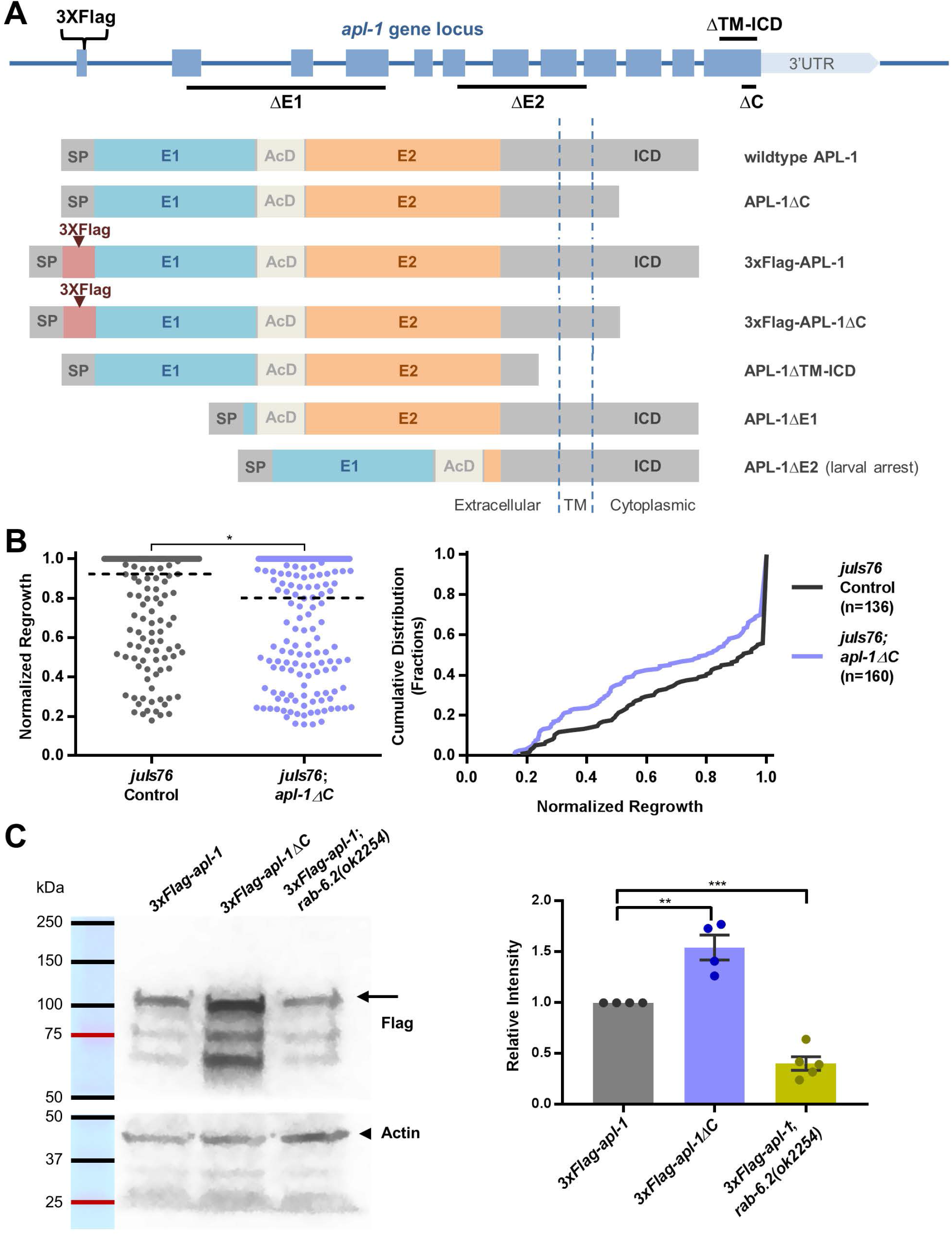
The cytoplasmic C-terminus of endogenous APL-1 is not required for the inhibition of axon regeneration in GABAergic motor neurons. (A) Diagrams showing CRISPR/Cas9 mediated gene editing of the endogenous *apl-1* locus, and the resulting protein produced. (SP signal peptide, AcD acidic domain, ICD intracellular domain, TM transmembrane domain, ΔC cytoplasmic c-terminal deletion) (B) Axon regeneration in *apl-1ΔC* mutant *(wp22)* is slightly decreased compared to *juIs76* control. (Kolmogorov-Smirnov D=0.1684; P=0.0309) (C) Western blot comparing 3xFlag tagged endogenous APL-1 protein level in control, *apl-1ΔC* mutant, and *rab-6.2(ok2254)* mutant. Arrow indicates full length 3xFlag-APL-1 band. 3xFlag-APL-1ΔC migrates at a slightly smaller size. Arrow head indicates actin (loading control). Quantification on the right shows four independent experiments (two lanes of *rab-6.2* mutant were loaded for one of the experiments therefore 5 data points total). One way ANOVA and Tukey’s multiple comparisons test is used. Column and error bar shows mean with SEM. 3xFlag-APL-1ΔC protein level is increased compared to 3xFlag-APL-1 control (Mean Difference=0.5425; P=0.0024). 3xFlag-APL-1 protein level is decreased in *rab-6.2(ok2254)* background compared to control (Mean Difference=-0.5989; P=0.0008).

Since RAB-6.2 is required for normal APL-1 levels (Figure 3A, 3B, 3C), and APL-1 itself inhibits axon regeneration, we predicted that *rab-6.2* null mutants would have increased regeneration. Indeed, we found that in *rab-6.2(ok2254)* null mutants, GABAergic axon regeneration was significantly increased (Figure 3D), indicating that the endogenous RAB-6.2 inhibits axon regeneration. Next, we asked whether *rab-6.2* inhibits axon regeneration specifically via its effect on APL-1. To address this question, we tested the genetic interaction between *rab-6.2* and *apl-1* in the inhibition of axon regeneration. We found that while GABAergic axon regeneration was increased by a greater degree in the *rab-6.2(ok2254)* null mutant than in the *apl-1* GABA-specific knockout mutant, the double mutant of the two did not increase regeneration further than the *rab-6.2* single mutant (Figure 3E). This suggests that while *apl-1* depends on *rab-6.2* to inhibit axon regeneration in the GABAergic motor neurons, it is unlikely the only axon regeneration inhibitor whose protein level or function depends on *rab-6.2.*

### The Cytoplasmic C-terminus of APL-1 is Not Required for the Inhibition of Axon Regeneration

To identify how APL-1 inhibits axon regeneration, we sought to identify the domains of APL-1 that are necessary for this novel function. The cytoplasmic tail of the APP protein family contains highly conserved regulatory motifs that serve as binding sites for adaptor proteins and have been shown to affect the trafficking and processing of the APP protein family (reviewed in Haass et al., 2012; Müller et al., 2017). The cytoplasmic domain can also be released into the cytoplasm following cleavage. This soluble fragment is known as AICD and has been implicated in transcriptional regulation (Cao and Südhof, 2001; Gao and Pimplikar, 2001; Kimberly et al., 2001; Konietzko, 2012; Müller and Zheng, 2012; Nhan et al., 2015). Thus, we asked if the cytoplasmic tail, including AICD, is necessary for APL-1’s inhibition of axon regeneration. To test this, we deleted most of the endogenous APL-1’s cytoplasmic tail via CRISPR/Cas9 mediated gene editing (APL-1ΔC), leaving the juxtamembrane residues intact for membrane anchoring (Figure 4A). Unlike deletion of the entire *apl-1* gene, which improved regeneration (Figure 2B), deletion of the cytoplasmic tail suppressed regeneration below wild type levels (Figure 4B). Thus, the C-terminal cytoplasmic tail of APL-1, including AICD, is not required for the inhibition of axon regeneration.

Why does deletion of the cytoplasmic tail result in *lower* regeneration than wild type? We hypothesized that this deletion affects trafficking and turnover of APL-1, resulting in protein accumulation and reducing regeneration. To test this possibility, we introduced a 3xFlag tag to the N-terminus of the endogenous APL-1ΔC just after the signal peptide domain via CRISPR/Cas9 mediated gene editing (Figure 4A). We detected significantly increased levels of 3xFlag-APL-1ΔC compared to the control full-length 3xFlag-APL-1 (Figure 4C). Consistent with our results, in APPΔC knock-in mice APPΔC has prolonged protein half-life and increased cell surface expression (Ring et al., 2007), possibly due to motifs in the cytoplasmic tail of APP family members that mediate endocytosis and turnover (Koo and Squazzo, 1994; Perez et al., 1999).

### Overexpressed Extracellular APL-1 Inhibits Axon Regeneration of the GABAergic Motor Neurons

In *C. elegans*, the GABAergic motor neurons along the body of the worm are positioned between the hypodermis and the hypodermal basal lamina (Figure 5A). Since *apl-1* is known to be expressed in the hypodermis of the worm (Hornsten et al., 2007; Wiese et al., 2010), we asked whether the hypodermal APL-1 has any contribution to the regulation of axon regeneration in the GABAergic motor neurons. To test this, we conditionally knocked out *apl-1* from the hypodermis by crossing our *apl-1(floxed)* allele into a strain expressing hypodermal-specific Cre recombinase, *tmIs1029* (Kage-Nakadai et al., 2014), under the *Pdpy-7* promoter. The occurrence of the excision event at the *apl-1* gene locus could be confirmed by single worm PCR (Figure S5A). We found that axon regeneration of the GABAergic motor neurons was not affected in the hypodermal-specific *apl-1* knockout mutant compared to the hypodermal Cre control strain (Figure 5B). This suggests that there is no endogenous extrinsic contribution from the hypodermal APL-1 to the inhibition of GABAergic axon regeneration.

**Figure 5.**
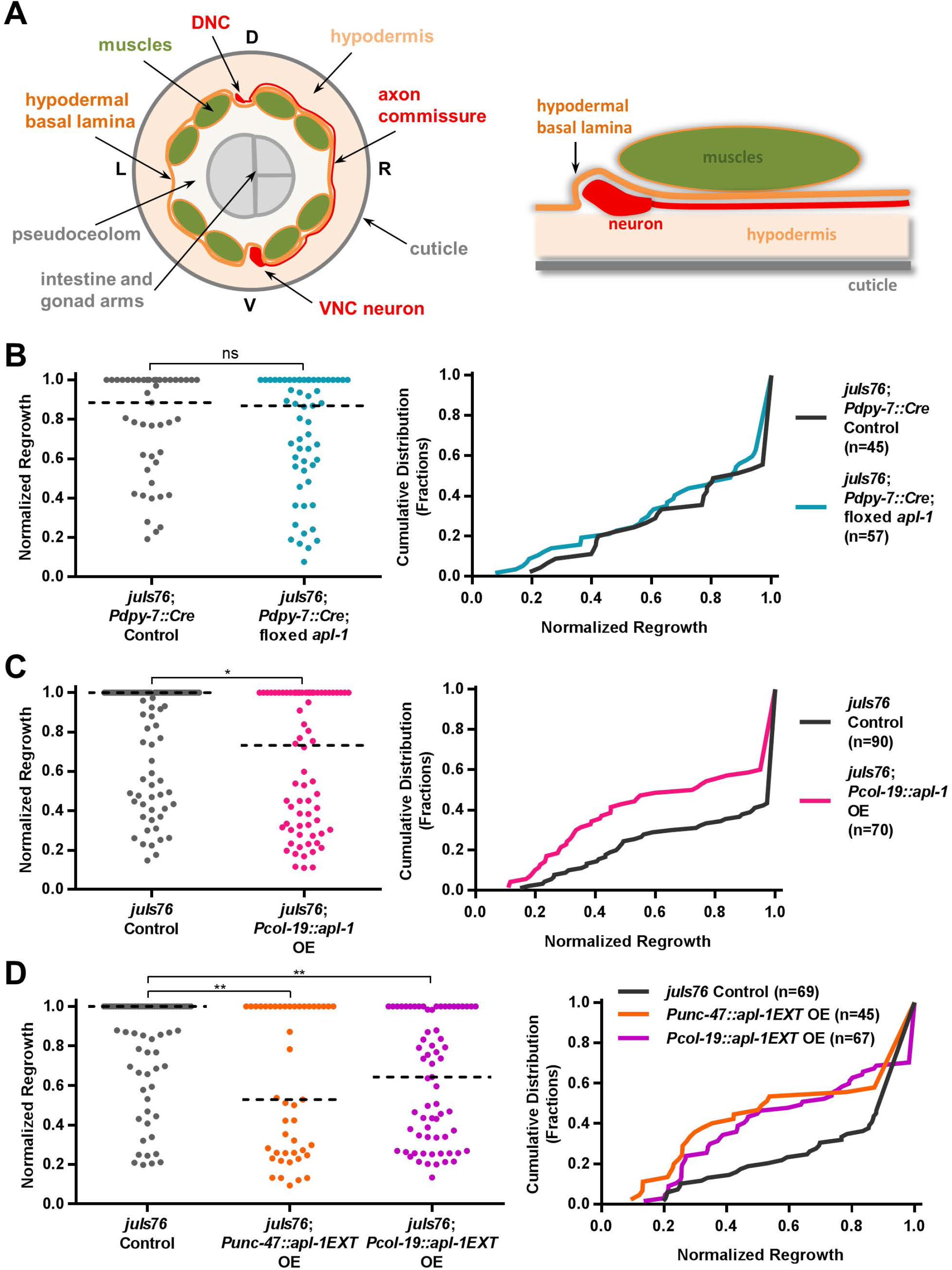
Secreted APL-1 ectodomain inhibits axon regeneration of GABAergic motor neurons. (A) Diagrams of a cross section of the worm showing the relative position of GABAergic neurons between the hypodermis and the hypodermal basal lamina. (not drawn to scale) (B) Axon regeneration is not changed in *apl-1* hypodermal cKO worms compared to hypodermal Cre only control background *(tmls1029).* (Kolmogorov-Smirnov D=0.1053; P=0.9433) (C) Axon regeneration is decreased in worms overexpressing *apl-1* specifically from the hypodermis *(col-19* promoter) compared to the *juls76* control. (Kolmogorov-Smirnov D=0.2254; P=0.0366) (D) Axon regeneration is decreased in worms overexpressing the APL-1 ectodomain fragment specifically from the GABAergic neurons *(unc-47* promoter) or from the hypodermis *(col-19* promoter) compared to the *juIs76* control. (GABA OE vs control Kolmogorov-Smirnov D=0.3304, P=0.0052; hyp OE vs control Kolmogorov-Smirnov D=0.2944, P=0.0055)

Lack of a phenotype in hypodermal apl-1-deleted animals could be because APL-1 expressed from the hypodermis cannot affect GABAergic motor neuron axon regeneration. Alternatively, the lack of phenotype could be because *apl-1* expression in the hypodermis is very low or insignificant after the L4 stage. It is known in *C. elegans* that *apl-1* expression in the hypodermal seam cells is repressed at the adult stage by the *let-7* microRNA (Niwa et al., 2008). To determine whether increased hypodermal APL-1 could affect regeneration, we overexpressed full-length APL-1 from two hypodermal promoters: *Pcol-19*, which is specific for the adult stage starting at the L4/adult molt (Liu et al., 1995), just after the time we perform axotomy at L4; and *Pdpy-7*, which is expressed only in the larval stages before the L4/adult molt (Gilleard et al., 1997; Johnstone and Barry, 1996). We found that overexpressing APL-1 from the *Pcol-19* promoter inhibited GABAergic axon regeneration (Figure 5C), while overexpressing APL-1 from the *Pdpy-7* promoter had no effect (Figure S5B). These data suggest that overexpressed cell-extrinsic APL-1 is able to inhibit GABAergic axon regeneration, and that it acts during the time the axons are regenerating. The lack of effect from earlier expression is consistent with the known fast turnover rate for the APP family of proteins (Lyckman et al., 1998; Morales-Corraliza et al., 2009; Perez et al., 1999). Further, these data indicate that while cell-extrinsic APL-1 from the hypodermis has the ability to inhibit the regenerative capacity of the GABAergic motor neurons, under normal conditions its expression levels are too low to do so.

### The Secreted Ectodomain of APL-1 Inhibits Axon Regeneration Independent of Endogenous GABAergic Neuronal Transmembrane APL-1

To further define the functional region of APL-1 that can inhibit regeneration, we tested whether the ectodomain of APL-1 is sufficient to suppress axon regeneration. We overexpressed the ectodomain fragment of APL-1 (APL-1EXT) (Figure 7B) specifically in the GABAergic neurons using the *Punc-47* promoter and also specifically from the hypodermis using the *Pcol-19* promoter. Since this domain contains a signal sequence but no transmembrane domain, overexpressed APL-1EXT fragment should be secreted into the extracellular region surrounding the neurons. We found that overexpressing APL-1 EXT in the GABAergic neurons or from the hypodermis both inhibited GABAergic axon regeneration (Figure 5D). These results are consistent with the decreased regeneration we observed in *apl-1 ΔC* mutants, since this truncated protein mainly consists of the ectodomain (Figure 4B).

The ectodomain of the APP protein family can dimerize with another APP family protein in *cis* or in *trans* (Soba et al., 2005; Wang et al., 2009), and the secreted APP fragment can act as a ligand of the full-length transmembrane APP (Milosch et al., 2014). Thus, we asked whether full-length APL-1 is required for the inhibition of axon regeneration by the APL-1 ectodomain, for example by acting as the receptor for the secreted APL-1 fragment. To test this possibility, we used CRISPR/Cas9 mediated gene editing to delete both the transmembrane and the cytoplasmic domain of the endogenous APL-1, eliminating its transmembrane form (APL-1ΔTM-iCd) (Figure 4A). Like *apl-1ΔC* animals, and in contrast to *apl-1* null mutants, *apl-1ΔTM-ICD* animals were viable and healthy, consistent with previous reports that the ectodomain is sufficient for viability (Hornsten et al., 2007). We examined axon regeneration in *apl-1ΔTM-ICD* mutants, and found that regeneration was even lower in these animals than in the *apl-1ΔC* mutants (Figure 6A). We conclude that the APL-1 ectodomain is sufficient to inhibit axon regeneration, and that neither transmembrane anchoring nor the intracellular domain is required.

**Figure 6.**
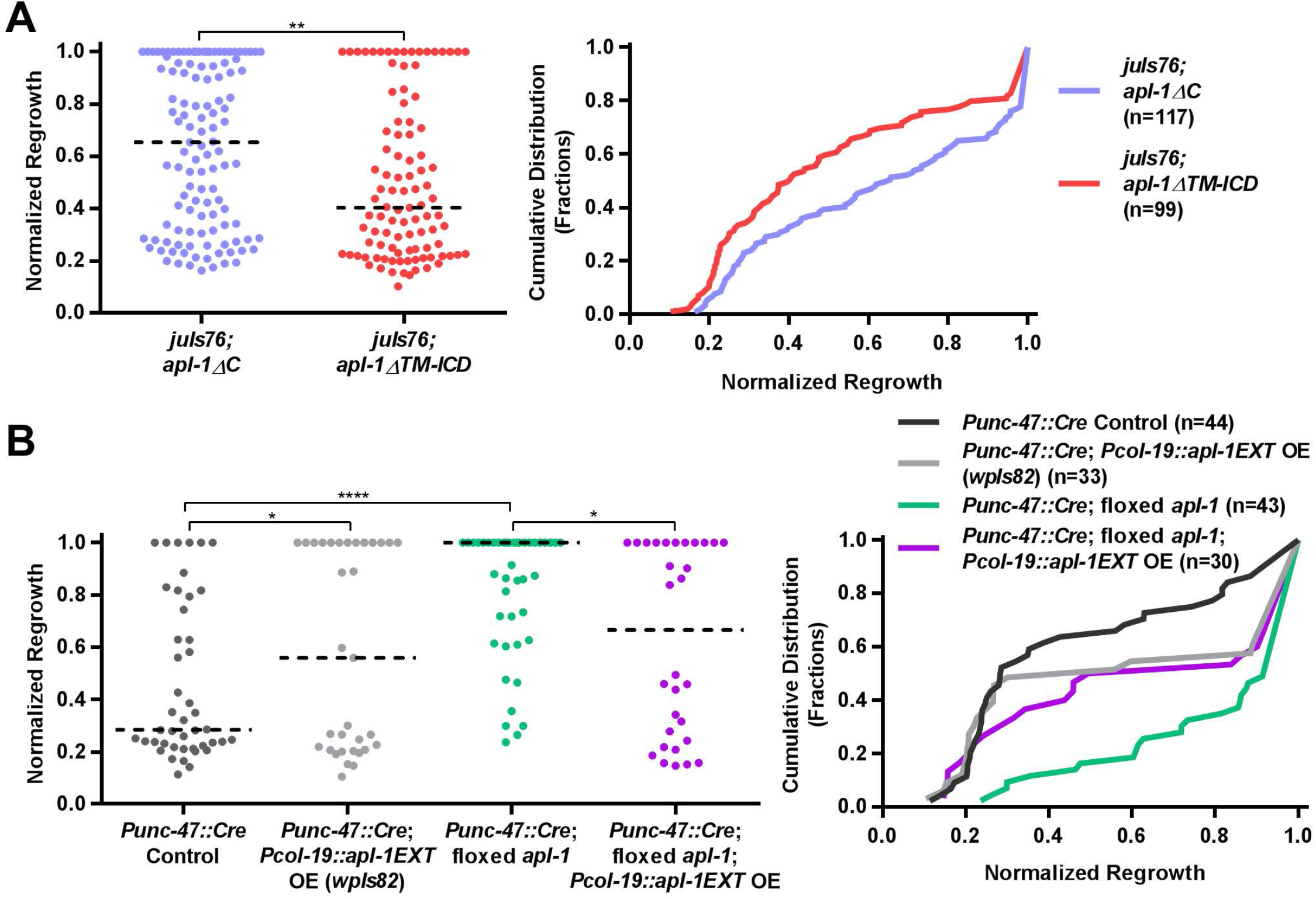
Secreted APL-1 ectodomain inhibits axon regeneration independent of endogenous neuronal APL-1. (A) Axon regeneration is lower in the *apl-1*Δ*TM-ICD* mutant than in the *apl-1ΔC* mutant (Kolmogorov-Smirnov D=0.2378; P=0.0047). (See Figure 4A for diagrams of mutant deletions.) (B) APL-1EXT overexpression from the hypodermis *(col-19* promoter) can still inhibit axon regeneration in the *apl-1* GABA-specific cKO background compared to the *apl-1* GABA specific cKO (Kolmogorov-Smirnov D=0.3504; P=0.0261). APL-1EXT overexpression increased regeneration in the Cre only control background which has lower baseline regeneration (Kolmogorov-Smirnov D=0.3182; P=0.0439). *(apl-1* GABA specific cKO vs Cre only control Kolmogorov-Smirnov D=0.5201, P<0.0001).

These data suggest that secreted APL-1 can inhibit axon regeneration even in neurons that completely lack full-length APL-1. We further tested this idea by examining the effect of secreted APL-1 expression in the *Pgaba*∷Cre; *apl-1(floxed)* background. To ensure consistent levels of APL-1EXT expression for comparison across different genetic backgrounds, we integrated the transgenic array overexpressing the secreted ectodomain from the hypodermis (*Pcol-19*∷APL-1EXT). To avoid potential effects from specific integrations, we tested two independent integrated lines. We found that on their own, both integrated lines overexpressing APL-1EXT inhibited GABAergic axon regeneration (Figure S5C), confirming our previous result using the extrachromosomal array (Figure 5D). We then crossed the integrated transgenes overexpressing APL-1EXT into the *Pgaba*∷Cre; *apl-1(floxed)* conditional knockout background. We found that APL-1EXT overexpression from the hypodermis inhibited GABAergic axon regeneration, even in the absence of the endogenous GABAergic neuronal APL-1 (Figure 6B, Figure S5D), similar to the effect of APL-1EXT overexpression in wild type animals (Figure S6C, 5D). Surprisingly, in the Pgaba∷Cre-only control background, APL-1EXT overexpression from the hypodermis increased regeneration (Figure 6B, S5D). We tested whether this could be due to non-specific effects of the transgene or the result of specific effects of the APL-1EXT protein, by overexpressing GFP under the same promoter *Pcol-19. Pcol-19*∷*GFP* overexpression did not increase regeneration in the Pgaba∷Cre controls (Figure S5E), suggesting that the effect on regeneration in the Pgaba∷Cre background is likely specific to the APL-1EXT protein, but the reason is not clear. Overall, however, these experiments, together with results from the *apl-1ΔTM-ICD* animals, indicate that inhibition of regeneration by APL-1EXT overexpression is not mediated by interactions with transmembrane neuronal APL-1.

The ectodomain of the APP protein family consists of an E1 domain and an E2 domain linked via a flexible acidic region. The E1 domain consists of a heparin binding and a copper binding subdomain and is important for APP dimerization (Dahms et al., 2010; Hoefgen et al., 2014; Kaden et al., 2008). To test if the E1 domain of endogenous APL-1 is involved in the inhibition of axon regeneration, we used CRISPR/Cas9 mediated gene editing to delete the E1 domain from the endogenous APL-1(Figure 4A). The resulting homozygous *apl-1ΔE1* mutant worms are viable without obvious phenotypes. We found that GABAergic axon regeneration was not affected in the *apl-1ΔE1* mutant worms (Figure 7A). We also tested the effect of overexpressing from the hypodermis APL-1EXT fragments with the E1 domain or the E2 domain deleted (*Pcol*-19∷APL-1EXTΔE1 and *Pcol*-19∷APL-1 EXTΔE2) (Figure 7B) and found that while APL-1EXTΔE1 overexpression can inhibit GABAergic axon regeneration, APL-1 EXTΔE2 overexpression did not have a significant effect (Figure 7C, 7D). Thus, the E1 domain is not responsible for the inhibition of GABAergic axon regeneration. These data indicate that the E2 domain is sufficient for APL-1’s function in regeneration.

**Figure 7.**
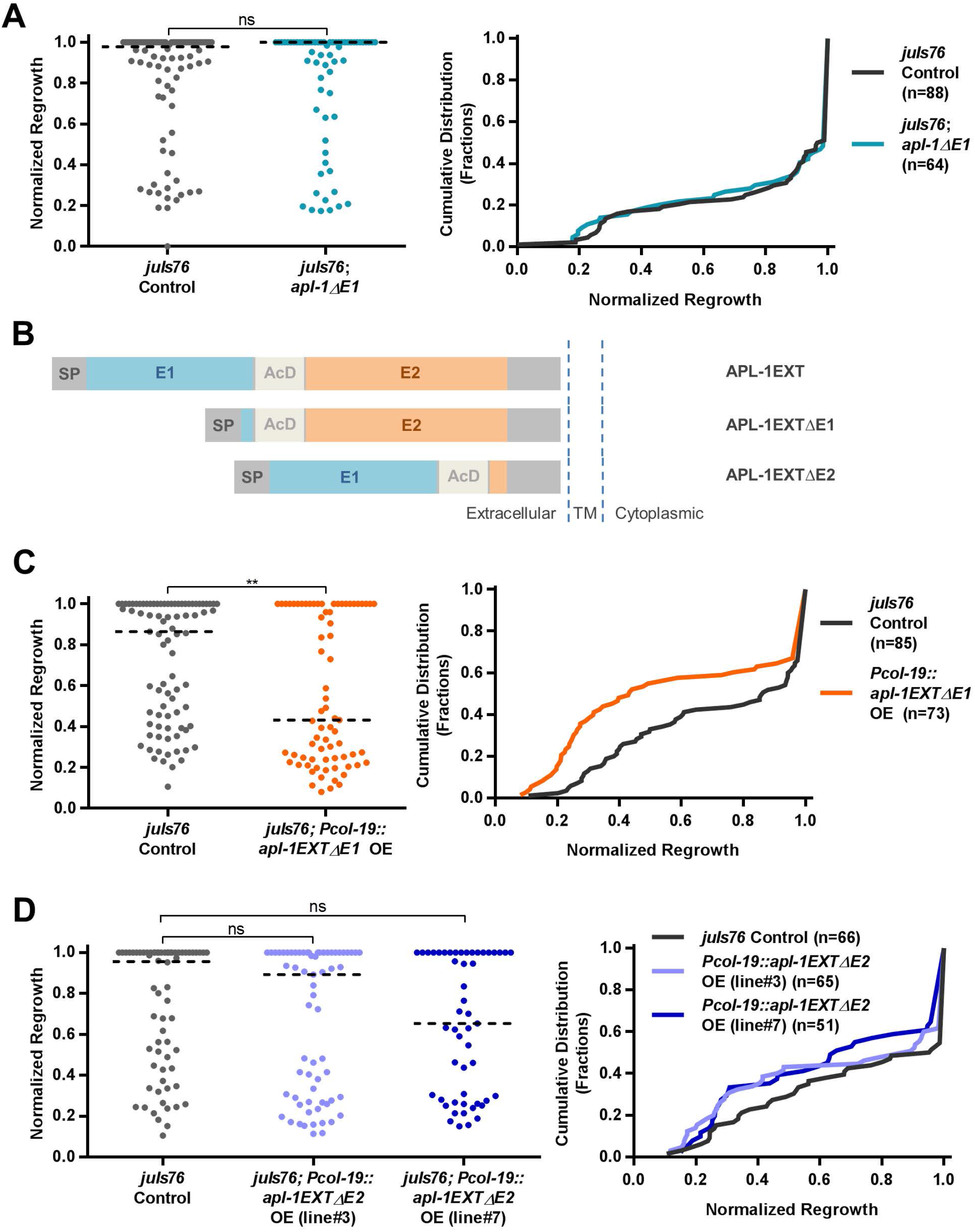
E2 domain of APL-1, but not E1 domain, regulates GABAergic axon regeneration. (A) Axon regeneration is not affected in the *apl-1ΔE1* mutant compared to the *juls76* control background (Kolmogorov-Smirnov D=0.06392; P=0.9981). See Figure 4A for diagrams of mutant deletions. (B) Diagram showing overexpression constructs of APL-1, including, APL-1EXT, APL-1EXTΔE1, and APL-1EXTΔE2. (C) APL-1EXTΔE1 overexpression is able to inhibit GABAergic axon regeneration in *juls76* control background (Kolmogorov-Smirnov D=0.2856; P=0.0033). (D) APL-1EXTΔE2 overexpression does not significantly affect GABAergic axon regeneration in *juls76* control background for two overexpression lines tested. (line#3 Kolmogorov-Smirnov D=0.1580, P=0.3867; line #7 Kolmogorov-Smirnov D=0.1818, P=0.2975)

### The Heparin Binding Site in the E2 Domain of the Endogenous APL-1 is Not Responsible for its Inhibition of Axon Regeneration

The E2 domain of the APP protein family contains binding sites for a number of extracellular matrix components including heparin, collagen, and F-spondin (Figure 8A) (Beher et al., 1996; Ho and Südhof, 2004; Hoopes et al., 2010; Lee et al., 2011; Wang and Ha, 2004; Xue et al., 2011). We used CRISPR/Cas9 mediated gene editing to delete the E2 domain from the endogenous APL-1, with the goal of using these animals to further explore the function of the E2 domain in regeneration. However, the resulting *apl-1ΔE2* mutant worms were not viable; they were arrested at L1/L2 similar to the *apl-1* null mutants. We next sought to create smaller mutations in the E2 domain, targeting conserved binding regions for ECM components. There are two heparin binding regions in the peptide sequence of E2 domain that form a single binding groove in the protein structure, and mutation of several residues in these two regions (N252A, H254A, R374A+K378A) has previously been shown to each reduce heparin binding affinity of APL-1 protein in vitro (Hoopes et al., 2010). We used CRISPR/Cas9 mediated gene editing to disrupt one of the heparin binding regions in the E2 domain of endogenous APL-1 by changing two conserved amino acids (R374A+K378A). These residues are also known to affect heparin binding affinity in human APP (Wang and Ha, 2004). The apl-1(R374A+K378A) mutant animals were viable without obvious phenotypes. However, axon regeneration was not affected in this mutant (Figure 8B). To attempt to more completely disrupt heparin binding, we used CRISPR/Cas9 mediated gene editing to mutate all four residues affecting heparin binding (N252A+H254A+R374A+K378A). Interestingly, these mutant animals were still viable, without obvious phenotypes, and no significant effect on axon regeneration was observed (Figure 8C).

**Figure 8.**
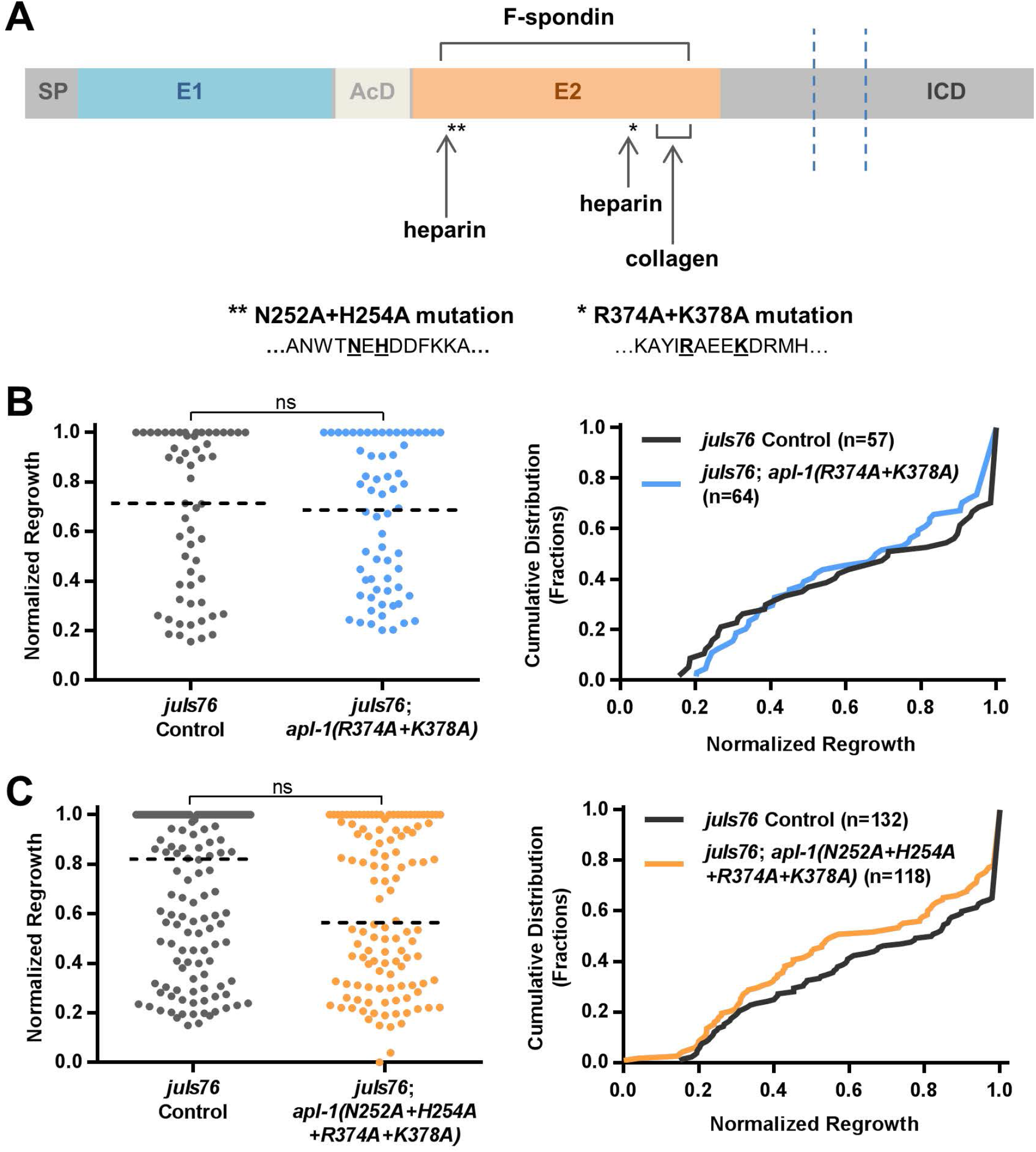
Heparin binding site in the E2 domain of endogenous APL-1 is not responsible for the inhibition of regeneration. (A) Diagram showing interaction sites in APL-1 E2 domain with different extracellular matrix components, and asterisks indicate the heparin binding site mutants tested. (B) Axon regeneration is not affected in the *apl-1(R374A+K378A)* mutant compared to *juls76* background (Kolmogorov-Smirnov D=0.1299; P=0.6888). (C) Axon regeneration is not significantly affected in the *apl-1(N252A+H254A+R374A+K378A)* mutant compared to the *juls76* background (Kolmogorov-Smirnov D=0.1439; P=0.1512). Mutating the heparin binding site does not abolish endogenous APL-1’s inhibition of regeneration.

Together, these data suggest that binding to heparin at the E2 domain is not required for APL-1’s function in either development or axon regeneration. Further experiments are required to identify the mechanism by which the E2 domain of APL-1 inhibits regeneration.

## Discussion

In this study, we investigated the role of APL-1 in regulating the regenerative capacity of GABAergic motor neurons after axonal injury in *C. elegans. C. elegans* offers an advantage for studying the function of the APP protein family without the complication of functional redundancies. We have found that the endogenous APL-1 inhibits axon regeneration in the GABAergic motor neurons. Our domain analysis data suggest that APL-1’s function in inhibiting axon regeneration does not require its cytoplasmic domain or its E1 domain. Rather, our data indicate that APL-1 inhibits axon regeneration through the function of its E2 domain. We have also shown that transmembrane anchoring is not required for APL-1 to inhibit axon regeneration. Together, our data indicate that APL-1 inhibits axon regeneration via its secreted E2 domain, independently of any other function of APL-1.

### ECM Components and Axon Regeneration

The E2 domain of APL-1 is highly conserved with other APP family proteins. APP and APL-1 are known to interact with a number of extracellular matrix (ECM) components via the E2 domain. There are two known heparin binding regions forming one binding groove, and also a collagen binding region in E2 (Beher et al., 1996; Hoopes et al., 2010; Lee et al., 2011; Wang and Ha, 2004; Xue et al., 2011). In addition, APP also binds F-spondin via its E2 domain (Ho and Südhof, 2004). The E2 domain has been shown to bind plasma membrane anchored heparan sulfate proteoglycans (HSPGs) and heparin chains (Dahms et al., 2015; Reinhard et al., 2013). One possible mechanism for APL-1’s inhibition of axon regeneration is via its interaction with certain ECM components, altering the adhesive properties of the neurons or changing the molecular nature of the surrounding ECM and how the extending axon responds. Many ECM molecules have been found to affect axon regeneration or neurite outgrowth. For example, the HSPG syndecan has been shown to regulate axon regeneration in *C. elegans* by promoting growth cone stabilization (Chen et al., 2011; Edwards and Hammarlund, 2014). Heparan sulfates have also been found to affect axon regeneration in mammalian peripheral nerves, although the exact role is unclear (Chau et al., 1999; Groves et al., 2005). The F-spondin *spon-1* mutant in C. elegans has been found to have increased axon regeneration (Chen et al., 2011), and while the *spon-1* mutant has largely normal motor neuron axon outgrowth in development, it synergistically enhances the outgrowth defects in the *unc-71* (an ADAM protein) mutant (Woo et al., 2008). It has been shown in rats that inhibition of collagen IV deposition at the injury site by antibodies or a drug that inhibits collagen triple helix formation can promote the regeneration of CNS neurons (Stichel et al., 1999). One of the type IV collagens in *C. elegans, emb-9*, has been found to be required for axon regeneration (Hisamoto et al., 2016). In vitro, collagen IV can promote axon outgrowth by functioning with integrin receptors (Bradshaw et al., 1995; Lein et al., 1991; Venstrom and Reichardt, 1995). Interestingly, it has been shown that the effects of APP knockout and sAPP overexpression on axon outgrowth of cultured neurons in vitro depend on β1 integrin (Young-Pearse et al., 2008), and there are data suggesting that APP and β1 integrin colocalize at the neuronal surface (Yamazaki et al., 1997) and they also interact biochemically (Young-Pearse et al., 2008). Thus, the secreted E2 domain of APL-1 may inhibit axon regeneration by interactions with ECM components.

### RAB-6.2 is a Multi-functional Inhibitor of Axon Regeneration

We found that APL-1 depends on RAB-6.2 to maintain its protein level, and that endogenous APL-1’s inhibition of axon regeneration is dependent on the presence of RAB-6.2. However, loss of *rab-6.2* increases axon regeneration more than the GABA-specific loss of *apl-1.* These data suggest that APL-1 is not the only axon regeneration inhibitor that depends on RAB-6.2. RAB-6.2 and its mammalian orthologue Rab6 are known to mediate the retrograde trafficking pathway from the endosome back to the TGN to recycle transmembrane proteins (Mallard et al., 2002; Young et al., 2005; Zhang et al., 2012, 2016), and Rab6 is a known regulator of APP trafficking via the adaptor protein X11/Mint (McConlogue et al., 1996; Thyrock et al., 2013). Thus, other direct or indirect effectors of RAB-6.2 likely also have a role in inhibiting axon regeneration, and might depend on functions of RAB-6.2 not limited to that in the retrograde recycling pathway. In *Drosophila* it has been found that the Rab6 ortholog Drab6 is a modulator of Notch signaling (Purcell and Artavanis-Tsakonas, 1999). Notch signaling in *C. elegans* have previously been found to inhibit axon regeneration (Bejjani and Hammarlund, 2012). It has also been shown in non-neuronal cells in both mammals and *C. elegans* that Rab6 is required for the retrograde trafficking and polarized relocalization of β1 integrin to facilitate cell migration (Shafaq-Zadah et al., 2016). RAB-6.2 function may thus be a critical choke point that can affect numerous independent mechanisms that inhibit axon regeneration, and targeting RAB-6.2 activity may be an effective way to improve regeneration. Interestingly, other Rab family members were recently found to be key inhibitors of axon regeneration in mouse and *C. elegans* (Sekine et al., 2018).

### APL-1 Trafficking and Processing Regulate Axon Regeneration

Our data indicate that the secreted APL-1 ectodomain inhibits axon regeneration. APP trafficking and processing are regulated by a complex network of adaptor proteins and interactors that typically bind to the conserved YENPTY motif in its cytoplasmic tail, and can be modulated by the TPEER motif upstream of the YENPTY motif as well (Müller et al., 2017). The adaptor protein X11/Mint, for example, binds the YENPTY motif to facilitate the regulation of APP trafficking by Rab6 and other regulators (Borg et al., 1996; McConlogue et al., 1996; Thyrock et al., 2013). Binding of the Pin1 prolyl isomerase to the phosphorylated threonine in the TPEER motif of APP can catalyze a conformational change in the proline, regulating APP processing likely as a result of alterations in trafficking (Pastorino et al., 2006). We found that axon regeneration in the *C. elegans* Pin1/*pinn-1* null mutant was significantly reduced (Figure S6A). However, Pin1 is also known to have other substrates and has been found to promote axon growth in cultured primary neurons by stabilizing CRMP2A (Balastik et al., 2015). We have also created more specific point mutations in the conserved TPEER motif of APL-1 by CRISPR/Cas9 mediated gene editing, including T to A and T to E mutations. We found that the APEER mutant had lower regeneration (Figure S6B), and the EPEER mutant had no effect on regeneration (Figure S6C). Together, these experiments do not clearly point to Pin1 as a key modulator of APL-1 in axon regeneration, and it remains unclear how these point mutations are affecting APL-1. In the broader perspective, our data predict that any manipulation that increases secretion of the APL-1 ectodomain will decrease regeneration, while reducing ectodomain secretion is likely to improve regeneration.

### APL-1 and Aldicarb Sensitivity

Aldicarb is an inhibitor of acetylcholinesterase which breaks down the excitatory neurotransmitter acetylcholine after its release by cholinergic neurons. Prolonged exposure of worms to aldicarb causes buildup of acetylcholine at the neuromuscular junctions (NMJ), leading to hypercontraction and eventually complete paralysis. Therefore, an aldicarb sensitivity assay can be used to measure synaptic function at the NMJ, which involves muscle innervation by both the excitatory cholinergic neurons (muscle contraction) and the inhibitory GABAergic neurons (muscle relaxation) (Miller et al., 1996; Vashlishan et al., 2008). We found that GABA-specific loss of *apl-1* leads to only slightly increased aldicarb resistance, an indication that there is increased GABA release, whereas *apl-1*Δ*TM-ICD* mutants are significantly hypersensitive to aldicarb and *apl-1*Δ*E1* mutants are similar to wildtype controls (Figure S2E, S2F). It has been previously reported that *apl-1* RNAi worms are hypersensitive to aldicarb (Wiese et al., 2010). While it is possible that RNAi treatment in worms have differential efficiencies on different neuronal cell types (Asikainen et al., 2005; Kamath et al., 2001; Timmons et al., 2001), their RNAi data and our data on the *apl-1*Δ*TM-ICD* and *apl-1*Δ*E1* mutants, which affect whole-body rather than GABA-specific APL-1, together appear to suggest that transmembrane APL-1 might play a role in inhibiting excitatory signal transmission as loss of transmembrane form of APL-1 in the *apl-1*Δ*TM-ICD* mutant and in the RNAi treated worms both led to aldicarb hypersensitivity. The greater effect size in these experiments compared to the small effect of GABA-specific knockout suggests that APL-1 might have a bigger role in cholinergic neurons in regulating synaptic transmission. Alternatively, and consistent with our results on regeneration, organism-wide reduction of *apl*-1 function might remodel the ECM and allow greater access of the drug to the NMJ, resulting in increased sensitivity. Further experiments would be needed to define the specific role of APL-1 in modulating aldicarb resistance.

### APL-1 and Neuronal Plasticity

The relation between axon regeneration and synapse formation has recently been discussed (Tedeschi et al., 2016). It has been pointed out that a number of genes that are required for proper function at the synapse are also inhibitors of axon regeneration, for example Tsc1, Pten, and Nf1 (Costa et al., 2002; Fraser et al., 2008; Goorden et al., 2007; Park et al., 2008; Romero et al., 2007), and studies have observed that synaptic-like differentiation or the presence of remaining synaptic branches post-injury can inhibit axon regeneration (Di Maio et al., 2011; Filous et al., 2014; Lorenzana et al., 2015; Wu et al., 2007). These data implicate the existence of genetic mechanisms that regulate the plasticity of neurons, controlling the balance between synaptic differentiation and axon growth potential. The regenerating axon needs to reach and innervate its target, and if it attempts synaptogenesis prematurely, its regrowth capacity could decline resulting in failed regeneration. Since APP has well known functions in promoting synaptic differentiation (Clarris et al., 1995; Dawson et al., 1999; Moya et al., 1994; Phinney et al., 1999; Schubert et al., 1991; Seabrook et al., 1999; Torroja et al., 1999; Wang et al., 2005, 2009; Yang et al., 2005) and we have found APL-1 inhibits axon regeneration, the APP protein family may act in general to regulate neuronal plasticity, favoring synaptic differentiation against sustained axonal regrowth.

## Material and Methods

### Strains

Worm strains were maintained at 20°C on NGM plates with 0P50 E. coli bacteria as the food source.

Strains containing the following alleles and transgenes were used:

*juIs76[Punc-25∷gfp + lin-15(+)]*

*wpIs40[Punc-47∷mCherry]*

*wpEx304[Papl-1∷mCherry]* 6.3kb promoter of apl-1

*wpEx305[Papl-1∷apl-1∷gfp]* 6.3kb promoter of apl-1

*wpEx223[Papl-1∷apl-1∷gfp; Peft-3∷Trap-2∷mCherry; Pmyo-2∷mCherry]* 5kb Papl-1

*wpEx301[Punc-47∷apl-1∷gfp; Punc-25∷mCherry; Pmyo-2∷mCherry]*

*apl-1(wp19)* floxed allele

*wpIs78[Punc-47∷nCre; Punc-47∷mCherry]*

*wpEx260[Punc-47∷nCre; Pmyo-2∷mCherry]*

*wpEx240[Punc-47∷apl-1; Pmyo-2∷mCherry]*

*wpEx289[Punc-47∷apl-1 ∷GFP∷SL2∷mCherry]*

*rab-6.2(ok2254)* obtained from CGC

*apl-1(wp29)* 3xFlag tagged allele

*wpEx303[Punc-47∷apl-1ΔC∷GFP∷SL2∷mCherry]*

*apl-1(wp22) apl-1ΔC*, deletion of C-term from TPEER to before stop codon

apl-1(wp30) 3XFlag-apl-lΔC

*tmIs1029[Pdpy-7∷nCre; Pgcy-10∷DsRed]* from Mitani Lab (Kage-Nakadai et al., 2014)

*wpEx239[Pdpy-7∷apl-1; Pmyo-2∷mcherry]*

*wpEx302[Pcol-19∷apl-1; Pmyo-2∷mcherry]*

*wpEx262[Punc-47∷apl-1EXT; Pmyo-2∷mCherry]*

*wpEx263[Pcol-19∷apl-1EXT; Pmyo-2∷mCherry]*

*wpIs81[Pcol-19∷apl-1EXT; Pmyo-2∷mCherry]* integration of *wpEx263*

*wpIs82[Pcol-19∷apl-1EXT; Pmyo-2∷mCherry]* integration of *wpEx263*

*apl-1(wp41) apl-1ΔTM-ICD*

*apl-1(wp34) apl-1ΔE1*

*wpEx296[Pcol-19∷GFP; pmyo-2∷mCherry]*

*wpEx298[Pcol-19∷apl-1EXTΔE1; Pmyo-2∷mCherry]*

*wpEx299[Pcol-19∷apl-1EXTΔE2; Pmyo-2∷mCherry]* (line#3)

*wpEx300[Pcol-19∷apl-1EXTΔE2; Pmyo-2∷mCherry]* (line#7)

*apl-1(wp35) apl-1(R374A+K378A)*

*apl-1(wp42) apl-1(N252A+H254A+R374A+K378A)*

*pinn-1(tm2235)* obtained from CGC

*apl-1(wp20)* APEER mutant

*apl-1(wp21)* EPEER mutant

**Table 1.**
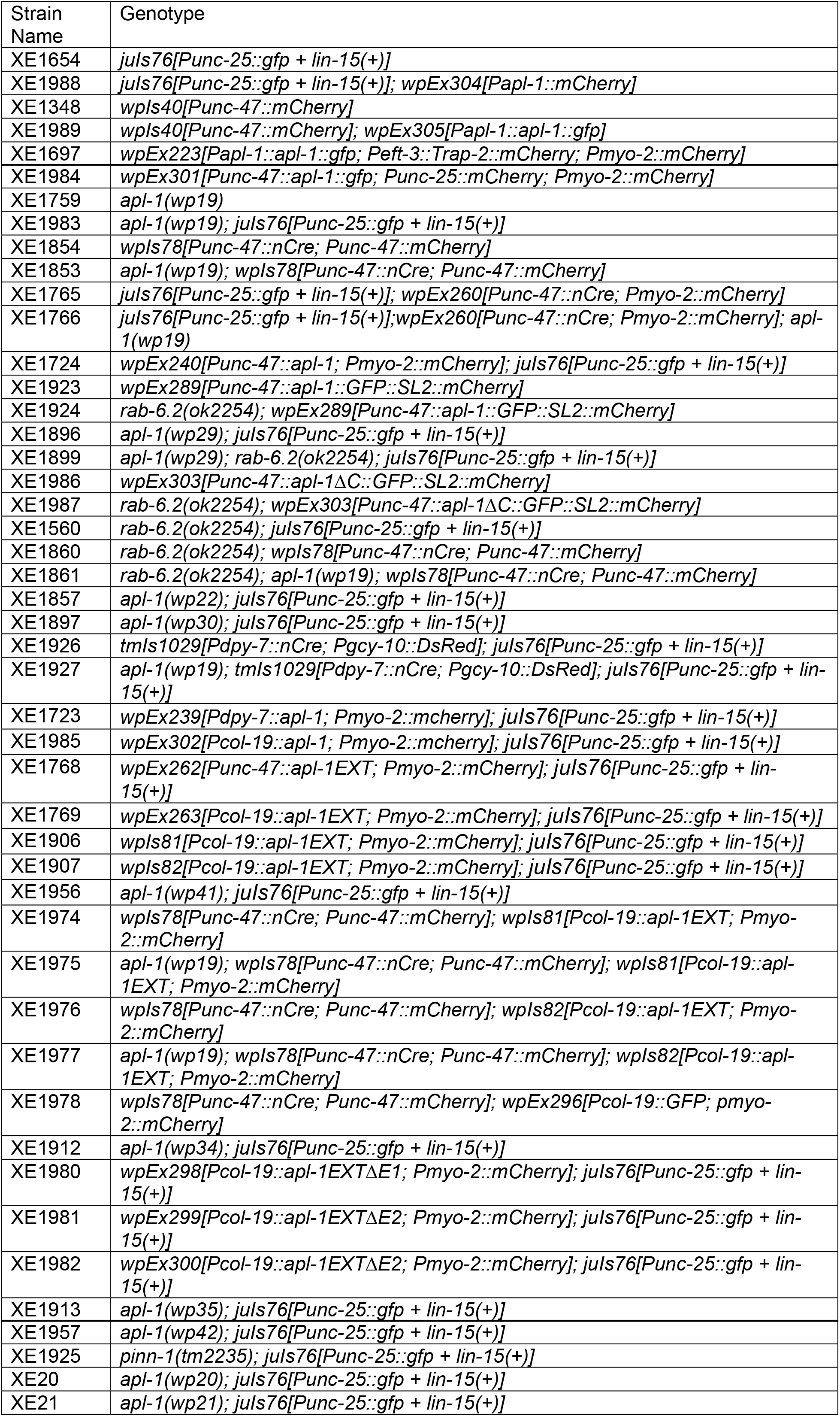
List of strains used.

### Transgene generation

Microinjection of plasmid DNA and TMP/UV integration of extrachromosomal arrays were performed according to standard procedures in worms (Mello et al., 1991).

### CRISPR/Cas9 mediated gene editing

*apl-1(wp19), apl-1(wp22), apl-1(wp29), apl-1(wp30), apl-1(wp34), apl-1(wp35), apl-1(wp20), apl-1(wp21), apl-1(wp41)*, and *apl-1(wp42)* were generated using CRISPR/Cas9 mediated gene editing technique with *dpy-10* co-conversion strategy to facilitate more efficient screening unless otherwise noted (method adapted from Arribere et al., 2014; Farboud and Meyer, 2015; Friedland et al., 2013; Paix et al., 2014). The Plasmids expressing Cas9 were obtained from Addgene #46168 (Friedland et al., 2013) and #47549 pDD162 (Dickinson et al., 2013) (p46168 was used initially but we later switched to pDD162 when creating *wp41* and *wp42*, which gave us higher edit efficiency). Plasmids expressing the desired sgRNAs were made using a preexisting sgRNA expressing plasmid as a PCR template, which was also obtained from Addgene #46169 (Friedland et al., 2013), swapping the targeting region with one of the PCR primers, and following with blunt end ligation of the PCR product. The first base of the 20bp targeting region is always changed to a G, resulting in a sequence of G(N)19, to enable expression under the U6 promoter. The *dpy-10* targeting sgRNA plasmid was made from the same template, using the published targeting region sequence GCTACCATAGGCACCACGAG (Arribere et al., 2014), and the repair template for converting *dpy-10* to the *cn64* dominant allele was ordered from IDT as an Ultramer oligo using the published oligo sequence AF-ZF-827 (Arribere et al., 2014). The repair templates for the editings in this paper are also single stranded oligos ordered from IDT with around 60bp homology recombination arms.

To do the CRISPR/Cas9 mediated gene editing, the following plasmid mix was injected into the gonad of young adult worms at the specified concentrations:

Peft-3∷cas9-SV40-NLS∷tbb-2 3’UTR at 50-66ng/ul

PU6∷dpy-10-sgRNA at 40-53ng/ul

dpy-10 repair template oligo AF-ZF-827 at 26ng/ul

PU6∷target-of-interest-sgRNA one or more at 50-66ng/ul each

Repair template oligo for target-of-interest at 70ng/ul

After recovering, injected P0 worms were singled to each plate and incubated at 25°C for 3 days. 40-50 P0’s were injected for each edit (only 20 P0’s when using pDD162 for expressing Cas9). The plate giving the biggest jackpot brood of *dpy-10* mutant worms was selected, and all the dumpy and/or roller F1 worms were individually isolated and allowed to lay eggs for 2 days before being lysed for PCR genotyping. For F1’s carrying the desired edit, their homozygous F2 progeny were isolated and confirmed with sequencing, and was outcrossed to remove the *dpy-10* edit.

The following sgRNA target sites, repair templates, and genotyping primers were used for each of the edits

#### apl-1(wp19)

5’ loxP insertion

(G)ATTAGTTTACCCACCGTCA (NGG)

(G)TTTTTTCAGCCATGACGGT (NGG)

cctcttttcttttcaacagattgacctatatttttttctaattcagacattttttcagccATAACTTCGTATAGCATACATTATACGAAGTTATatgacggtgggtaaactaatgattggcttacttataccgattcttgtcgccacagtttacgc

Genotyping Forward: gtgatgctacacagccatttc

Genotyping Reverse: gcaattaaacaaacatacctctgcg

3’ loxP insertion

(G)AAAAAAAGTACGGTCTTGG (NGG)

(G)AGTAAAAAAAGTACGGTCT (NGG)

(G)GTAGTGAGTAAAAAAAGTA (NGG)

caatggctatgaaaacccgacgtactcattcttcgactcgaaggcctaaaccaccaagacATAACTTCGTATAGCATACATTATACGAAGTTATcgtactttttttactcactactaccaacaattttacaccaaaatgatccattctcctatc

Genotyping Forward: cgtccagcgtcttccaaccatac

Genotyping Reverse: atacgaacaacaagcatgagtg

Note: the *dpy-10* co-conversion strategy was used in all CRISPR/Cas9 mediated gene editing experiments except for the *5’ loxP* site in the floxed *apl-1* allele *wp19. apl-1(wp19)* was generated by first inserting the *5’ loxP* site to obtain *apl-1(wp17)* and then inserting the 3’ loxP site on top of it. *apl-1(wp17)* was created before the usage of the *dpy-10* co-conversion method. Only the Cas9 plasmid, the specific targeting sgRNA’s and the repair template were injected along with a fluorescent co-injection marker *Pmyo-2∷mCherry*, and the selection of potential edited F1 worms for genotyping was based on the expression of the fluorescent co-injection marker *Pmyo-2∷mCherry*.

#### apl-1(wp22)

(G)GTAGACGTCTACACACCAG (NGG)

(G)AGTACGGTCTTGGTGGTTT (NGG)

gccatcaccaacgctcgtcgtcgccgtgccatgcgcggtttcatcgaggtagacgtctac-

taaaccaccaagaccgtactttttttactcactactaccaacaattttacaccaaaatgatcca

Genotyping Forward: cgtccagcgtcttccaaccatac

Genotyping Reverse: atacgaacaacaagcatgagtg

#### apl-1(wp29)

(G)GTCGCCACAGTTTACGCAG (NGG)

gacggtgggtaaactaatgattggcttacttataccgattcttgtcgccacagtttacgcaGACTACAAAGACCATGACGGTGATTATAAAGATCATGACATCGATTACAAGGATGACGATGACAAGgaggtatgtttgtttaattgcaaaggggtttgcgcgggtcgattcccactgactcgtaac

Genotyping Forward: gtgatgctacacagccatttc

Genotyping Reverse: gaacatctagtcgagagttctagtc

#### apl-1(wp30)

was created by editing on top of *apl-1(wp22)* using same sgRNA and repair template as for making *apl-1(wp29).*

#### apl-1(wp34)

(G)GTCTTCCATGATCCCTCTT (NGG)

(G)ATGGTAACTTACGGTCATT (NGG)

atatcatcttatgacacttacaactgactcatctggaaactgtcatggtaacttacGGTCATTCTCTTCGGTCATATACTG

GTTGCGGTATCCACATGAAAATGCGACCATTGGAATGAACTTCT

Genotyping Forward: gtgatgctacacagccatttc

Genotyping Reverse: TAGTCCTTCCAGTAGGTGACTGC

#### apl-1(wp35)

(G)TACATCCGTGCAGAGGAGA (NGG)

TACTCACGTCAACAAGCCAAACAAGCACTCTGTTCTCCAATCTCTTAAGGCTTACATCgcTG

CAGAGGAGgccGATCGCATGCACACTTTGAACAGATACCGTCACTTGCTGAAGGCCGATTC

AAAGGAAGCT

Genotyping Forward: GGAGATTTGGAGACGAGATAC

Genotyping Reverse: TAGTCCTTCCAGTAGGTGACTGC

#### apl-1(wp20)

(G)GTAGACGTCTACACACCAG (NGG)

TGCCATCACCAACGCTCGTCGTCGCCGTGCCATGCGCGGTTTCATCGAGGTAGACGTCTA

CgctCCAGAGGAGCGTCATGTCGCTGGAATGCAAGTCAATGGCTATGAAAACCCGACGTAC

TCA

Genotyping Forward: CGTCCAGCGTCTTCCAACCATAC

Genotyping Reverse: atacgaacaacaagcatgagtg

#### apl-1(wp21)

(G)GTAGACGTCTACACACCAG (NGG)

TGCCATCACCAACGCTCGTCGTCGCCGTGCCATGCGCGGTTTCATCGAGGTAGACGTCTA

CgaaCCAGAGGAGCGTCATGTCGCTGGAATGCAAGTCAATGGCTATGAAAACCCGACGTAC

TCA

Genotyping Forward: CGTCCAGCGTCTTCCAACCATAC

Genotyping Reverse: atacgaacaacaagcatgagtg

#### apl-1(wp41)

(G)TGATTGGATGAGCTTGTCG (NGG)

(G)AGTACGGTCTTGGTGGTTT (NGG)

Repair 1:
AGAGCTTCGTGTCGACATTGAGCCGATCATCGATGAGCCAGCCTCATTCTACCGCCACGACAAGTAAaccaccaagaccgtactttttttactcactactaccaacaattttacaccaaaatgatcca

Repair 2:
ggatcattttggtgtaaaattgttggtagtagtgagtaaaaaaagtacggtcttggtggtTTACTTGTCGTGGCGGTAGAATGAGGCTGGCTCATCGATGATCGGCTCAATGTCGACACGAAGCTCT

Note: both repair template oligos were used.

Genotyping Forward: CAATAGAAGAGGAGAAGAAGGCTCC

Genotyping Reverse: atacgaacaacaagcatgagtg

#### apl-1(wp42)

was created by editing on top of *apl-1(wp35)*

(G)TCGTGCTCGTTGGTCCAGT (NGG)

(G) TTAAAATCGTCGTGCTCGT (NGG)

Repair 1:
CTTGTCAACCTTCTTTCTGTGCTTCTCATCCATTCTCATTTCTGCCTTCTTAAAATCGTCagcCTCtgcGGTCCAGTTGGCAATCTTGAAGTATGGATCTTGGGAACTTGGTTCCTCTTCGTCCTTCTCGTCGG

Repair 2:
CCGACGAGAAGGACGAAGAGGAACCAAGTTCCCAAGATCCATACTTCAAGATTGCCAACTGGACCgcaGAGgctGACGATTTTAAGAAGGCAGAAATGAGAATGGATGAGAAGCACAGAAAGAAGGTTGACAAG

Note: both repair template oligos were used.

Genotyping Forward: TTTGCCGTTCTTGAGCCATGC

Genotyping Reverse: ACTTCTCGGCTCCCTTTGGATC

### Western blots

300 mid/late L4 worms were collected in 30ul M9 with 0.1% Tween20 and mixed with 30ul 2x Laemmli sample buffer with β-Mercaptoethanol (Bio-Rad), freeze-thawed 4 times to rupture the cuticle, denatured at 70°C for 20 minutes, and loaded onto 7.5% precast polyacrylamide gels (Bio-Rad). Gels were run at 125V for 45 minutes and were transferred to PVDF membrane overnight (14-16 hours) at 4C at 30-50mA. The membrane was cut in half at 50 kD, and the top half was probed with mouse anti-Flag antibody (Sigma F3165, 1:5000), whereas the bottom half was probed with mouse anti-β-actin antibody C4 (Santa Cruz sc-47778 1:2500). TBST was used for washes and 5% milk in TBST was used for blocking. Anti-mouse HRP secondary antibody, Super-Digital ECL solution, and the KwikQuant Imager from Kindle Biosciences LLC were used. Protein band intensities were measured in ImageJ, normalized to background intensities, and the 3xFlag-APL-1 protein levels were normalized to actin.

### Fluorescent Imaging and Quantification

Imaging for scoring axon regeneration, time lapse imaging of APL-1-GFP trafficking, and imaging for APL-1-GFP and Trap-2-mCherry colocalization were done with Olympus BX61 microscope with attached Olympus DSU, Olympus UPlanFL N 40X/1.30 oil objective or UPlanSApo 60X/1.35 oil objective, and either Andor Neo sCMOS camera or Hamamatsu C11440 camera. Worms were mounted on 3% agarose gel pads and immobilized in 0.05 um polystyrene microbeads or 0.1% levamisole in M9.

The other images for APL-1 expression and localization were done with an UltraView Vox (PerkinElmer) spinning disc confocal microscope (Nikon Ti-E) and Hamamatsu C9100-50 camera using 40X CFI Plan Apo, NA 1.0 oil objective and Volocity software (Improvision). 0.1% levamisole in M9 was used to immobilize the worms.

For the quantification of the expression level of Punc-47∷APL-1-GFP∷SL2∷mCherry in N2 versus *rab-6.2(ok2254)*, the GFP channel was set with a 488nm laser power of 15%, sensitivity 100, exposure 50ms, and the RFP channel was set with a 561nm laser power of 24%, sensitivity 100, exposure 50ms consistently. Images were taken with 0.1um Z-step. Image stacks were exported to ImageJ in 16bit format, and a SUM of slices Z projection was applied to the stack. Quantification was done by selecting an oval ROI encompassing the cell body, and then normalizing the integrated intensity of the green channel to the red channel. The distribution of the normalized expression of APL-1-GFP was compared between N2 and *rab-6.2(ok2254)* mutant using the non-parametric Mann-Whitney U test. And for the quantification of the expression level of Punc-47∷APL-lΔC-GFP∷SL2∷mCherry in N2 versus *rab-6.2(ok2254)*, the GFP channel was set with a 488nm laser power of 12%, sensitivity 100, exposure 50ms, and the RFP channel with a 561nm laser power of 24%, sensitivity 100, exposure 50ms.

### Axotomy

Laser axotomy experiments were performed as previously described (Byrne et al., 2011). Axon regrowth was scored as the vertical distance between the growth cone end (or the injured axon stump end) and the ventral nerve cord, normalized to the corresponding vertical distance between the dorsal and ventral cords. As a result, full regeneration to the dorsal nerve cord is represented by a value of 1. The cumulative distribution of the normalized axon regrowth measurements was compared between control and experimental groups using the non-parametric Kolmogorov-Smirnov test.

### Aldicarb Assay

The day before the assay, aldicarb (Ultra Scientific PST940) was dissolved to obtain fresh 400μM stock in ethanol, and then diluted to 40μM in water. 40μM aldicarb solution was top-spread to NGM plates to obtain a final concentration of 1mM. Plates were allowed to dry and were then seeded with 100μL overnight culture of OP50 bacteria and incubated at room temperature overnight. On the day of the assay, 25 one-day young adult worms were placed on each plate and the fraction of worms paralyzed was counted every 20 minutes for two hours. Paralysis was defined as inability to respond to a touch stimulus. 3 trials were performed for each genotype. Two-way repeated measures ANOVA with Bonferroni’s multiple comparisons test was used to compare different strains over the time course.

---

**Figure S1.**
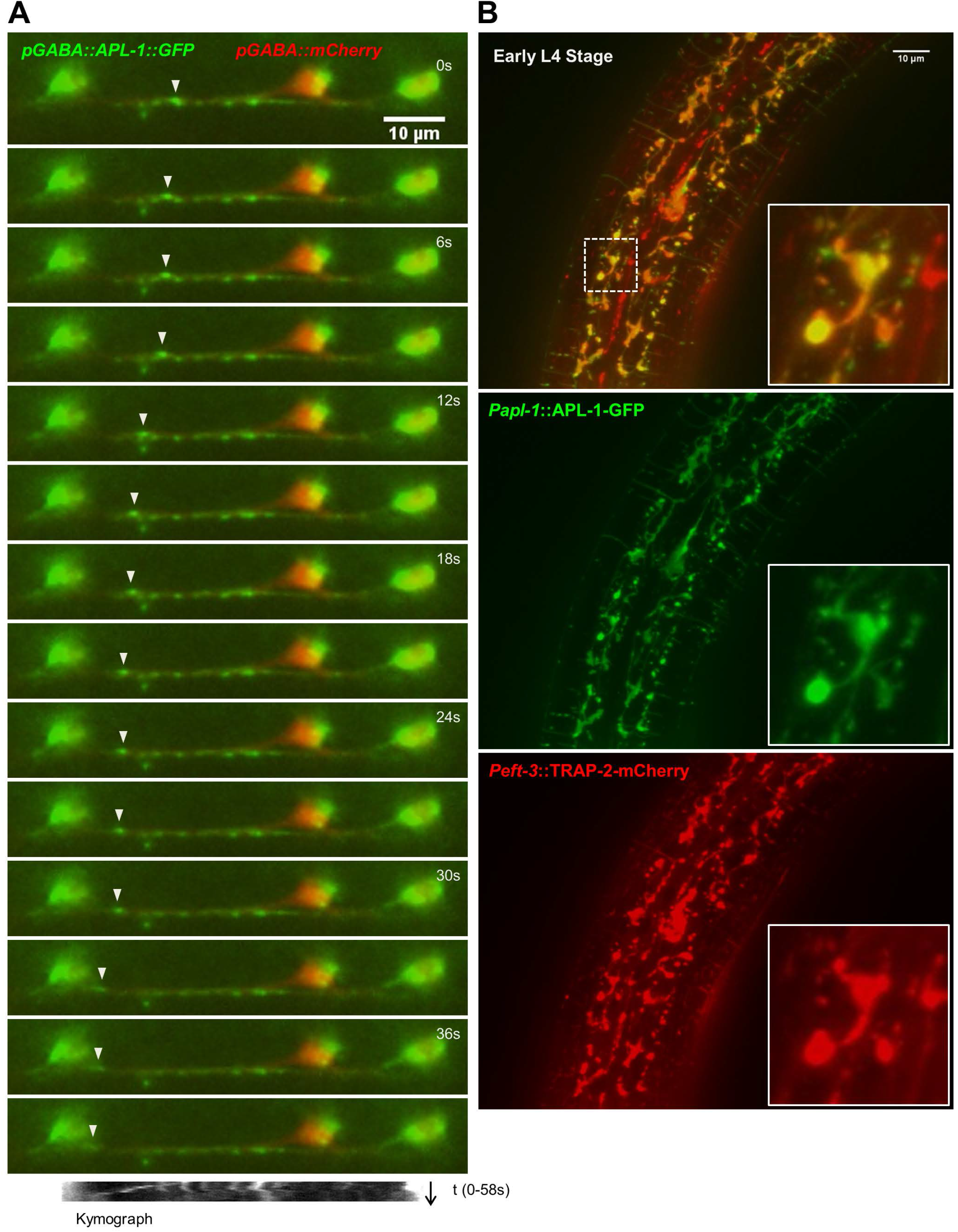
Subcellular localization and trafficking of APL-1-GFP (A) APL-1-GFP containing puncta (arrowhead) trafficking along axons in GABA neurons. Time lapse images each separated by 3 seconds. Scale bar 10μm. (B) *Papl-1*∷APL-1GFP expression found in subcellular compartments in the hypodermis of early L4 stage worms. The subcellular localization of APL-1-GFP co-localizes with an ER marker TRAP-2-mCherry and is also found in other vesicular compartments. Scale bar 10um. Inset shows enlargement of boxed area.

**Figure S2.**
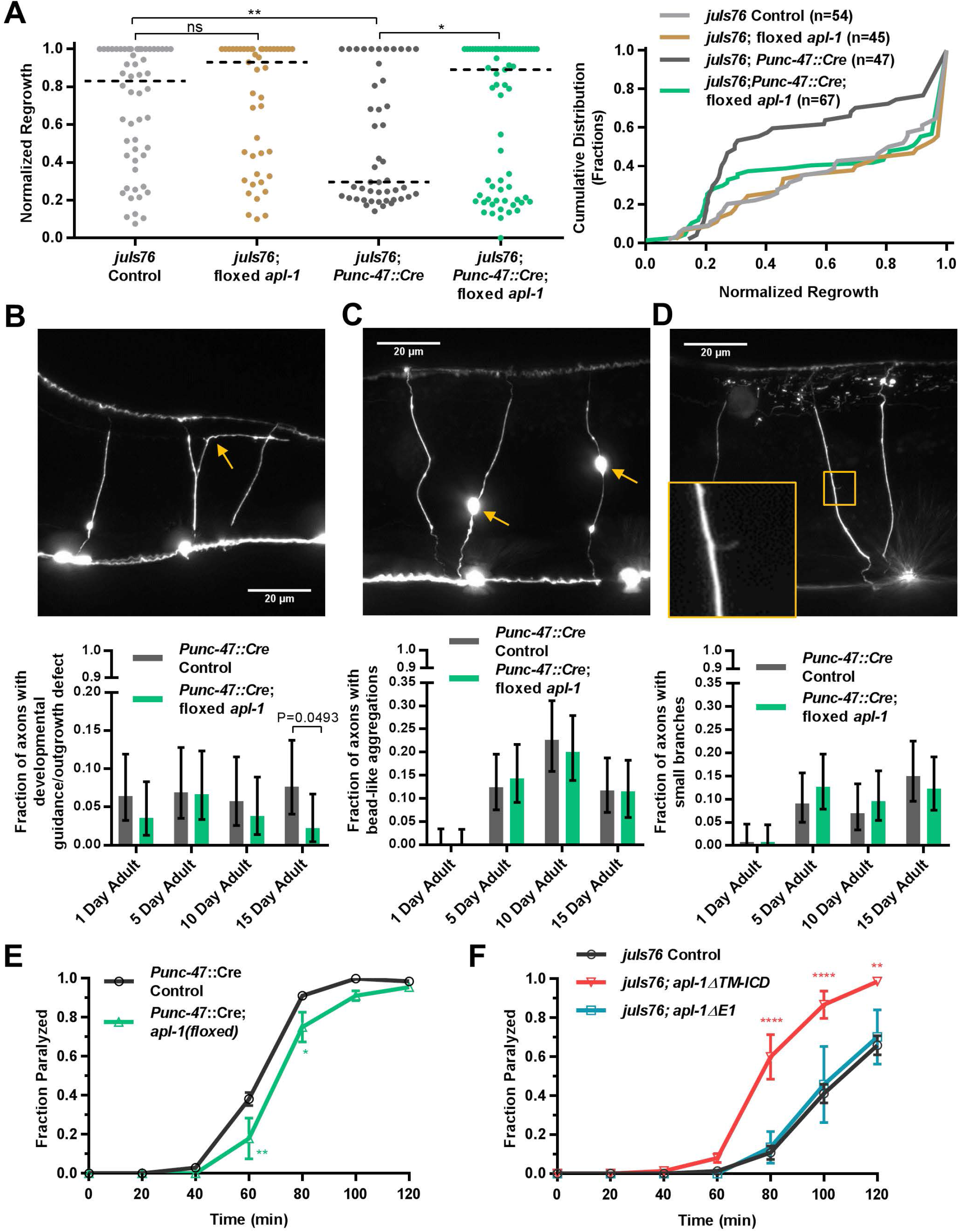
*apl-1* in axon morphology and function. (A) Axon regeneration is not affected by insertion of the *loxP* sites at the genomic locus (Kolmogorov-Smirnov D=0.1296; P=0.8039). Expression of Cre alone in GABA neurons *(unc-47* promoter, extrachromosomal array) lowers baseline regeneration compared to *juIs76* control (Kolmogorov-Smirnov D=0.3550; P=0.0036). GABA specific Cre-loxP *apl-1* cKO increases regeneration compared to Cre only control (Kolmogorov-Smirnov D=0.2991; P=0.0143). (B) Plot shows the fraction of GABA axons with developmental guidance or outgrowth defects in Cre only control versus GABA specific Cre-loxP *apl-1* cKO worms. Error bar shows 95% confidence interval. The fractions remain largely unchanged as the worm ages is consistent with the effect being developmental. Such defects are absent from wildtype worms (data not shown). Picture shows an example of an misguided axon (arrow). Scale bar 20um. (C) Plot shows the fraction of GABA axons with bead-like aggregations in Cre only control versus GABA specific Cre-loxP *apl-1* cKO worms. Error bar shows 95% confidence interval. This phenotype correlates with age. The decrease seen in 15 day old adults might be due to earlier death of worms that have more aggregations. Picture shows examples of bead-like aggregations (arrows). Scale bar 20um. (D) Plot shows the fraction of GABA axons with small branch-like protrusions Cre only control versus GABA specific Cre-loxP *apl-1* cKO worms. Error bar shows 95% confidence interval. Picture on the right shows an example of a small branch-like protrusion. Scale bar 20um. (E) Aldicarb sensitivity assay over a 120-minute time course comparing the fraction of worms paralyzed in *Punc-47*∷Cre controls versus *Punc-47*∷Cre; *apl-1(floxed)*, scored every 20 minutes. *Punc-47*∷Cre; *apl-1(floxed)* worms have slightly reduced sensitivity. Two-way repeated measures ANOVA with Bonferroni’s multiple comparisons test is used; P=0.0041 at 60min, P=0.0311 at 80min. Error bar shows SEM. (F) Aldicarb sensitivity assay over a 120-minute time course comparing the fraction of worms paralyzed in *juIs76* controls versus *apl-1*Δ*TM-ICD* mutants and *apl-1*Δ*E1* mutants, scored every 20 minutes. *apl-1*Δ*TM-ICD* mutants have significantly increased sensitivity compared to controls, whereas *apl-1*Δ*E1* mutants are comparable to controls. Two-way repeated measures ANOVA with Bonferroni’s multiple comparisons test is used; for *apl-1*Δ*TM-ICD* versus control P<0.0001 at 80min, P<0.0001 at 100min, P=0.0019 at 120min. Error bar shows SEM.

**Figure S3.**
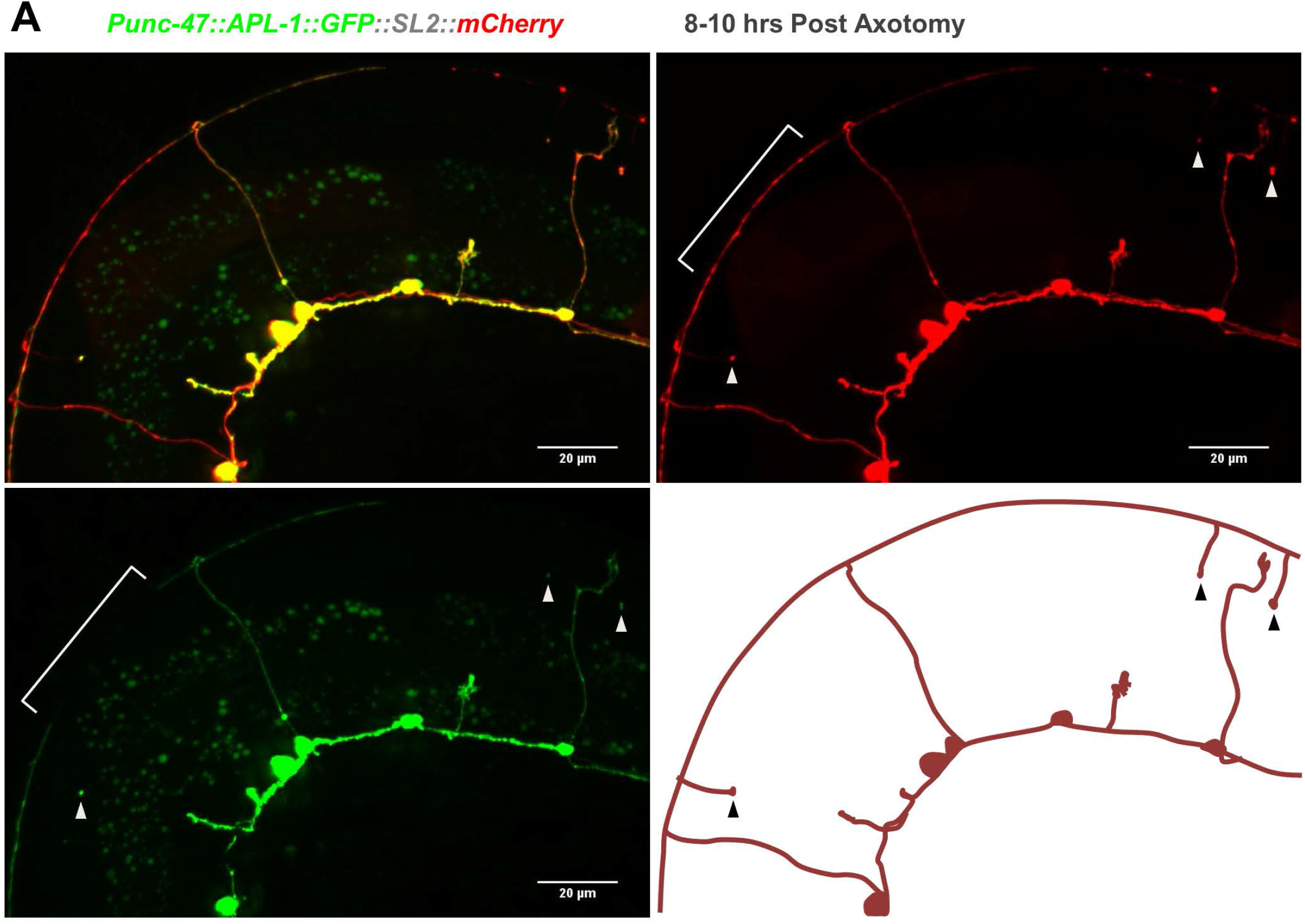
APL-1-GFP localization after injury. (A) APL-1-GFP localization 8-10 hours after axotomy. Proximal to the cut site, protein is seen in the growth cone or at the proximal tip, along the axons, and in the cell body. Distal to the cut site, accumulation is seen at the tip of the distal stump (arrowheads), but absent from the more distal segment (bracket). Scale bar 20μm. Lower right panel shows a diagram of the mCherry channel.

**Figure S4.**
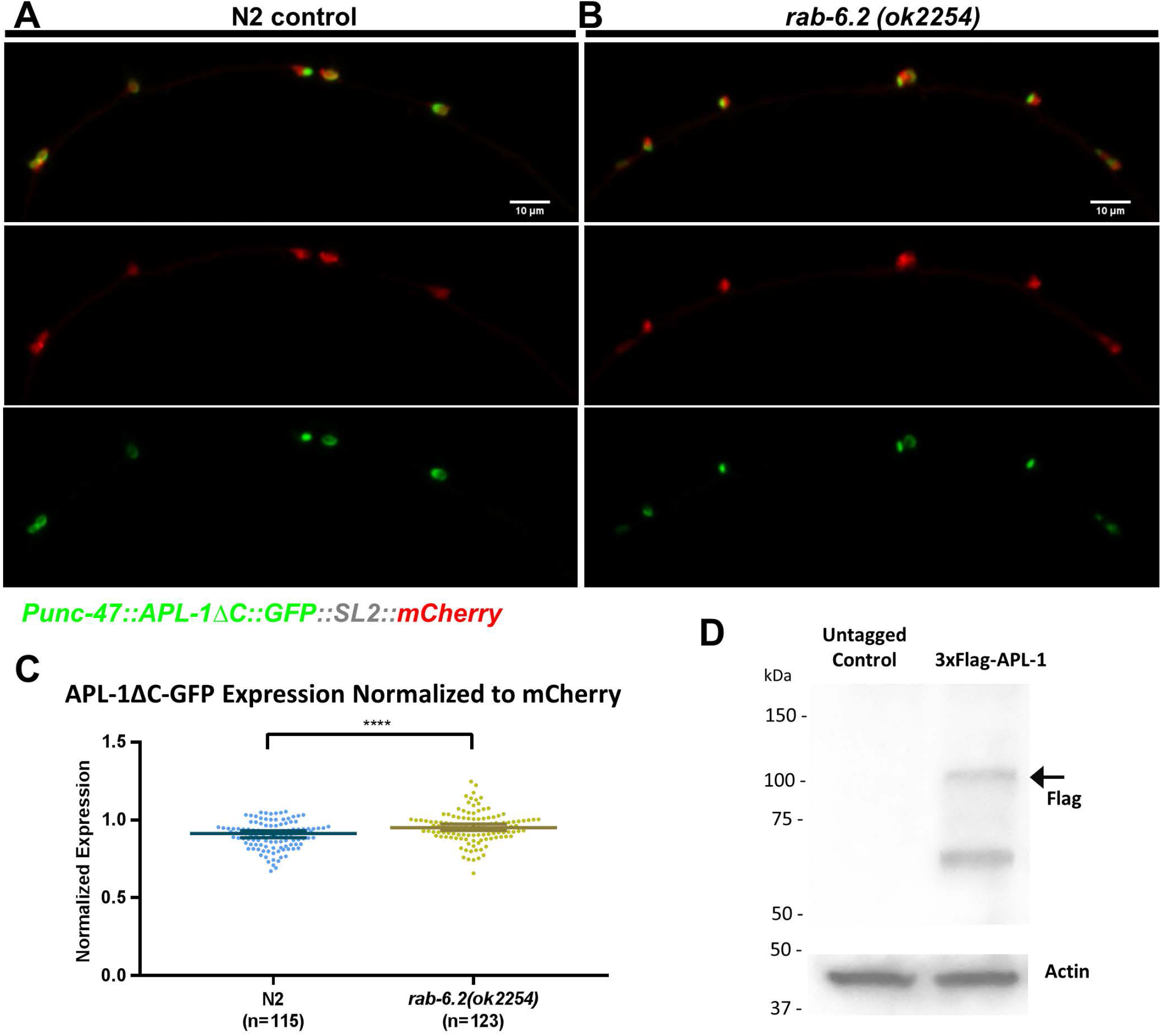
APL-1ΔC-GFP expression is not decreased in *rab-6.2(ok2254)* mutants. (A) APL-1ΔC-GFP expressed in the GABA neurons of N2 control worms, along with mCherry co-expressioin using SL2. Scale bar is 10μm. (B) Same expression as (A) in *rab-6.2(ok2254)* mutant worms. Scale bar is 10μm. (C) Quantification of APL-1ΔC-GFP expression in cell bodies normalized to mCherry, comparing N2 control (A) versus *rab-6.2(ok2254)* mutant (B). Mann-Whitney U test is used (control median=0.9129; *rab-6.2* median=0.9512; P<0.0001). Line and error bar shows median and 95% CI. The APL-1ΔC-GFP expression is not decreased in *rab-6.2(ok2254)*, and the slight increase is tiny in effect size. (D) Western blot using anti-Flag antibody against 3xFlag-APL-1 tagged worm versus untagged worm. Actin is used as loading control. The presence of the band corresponding to full length 3XFlag-APL-1 (arrow) is specific to the tagged worm.

**Figure S5.**
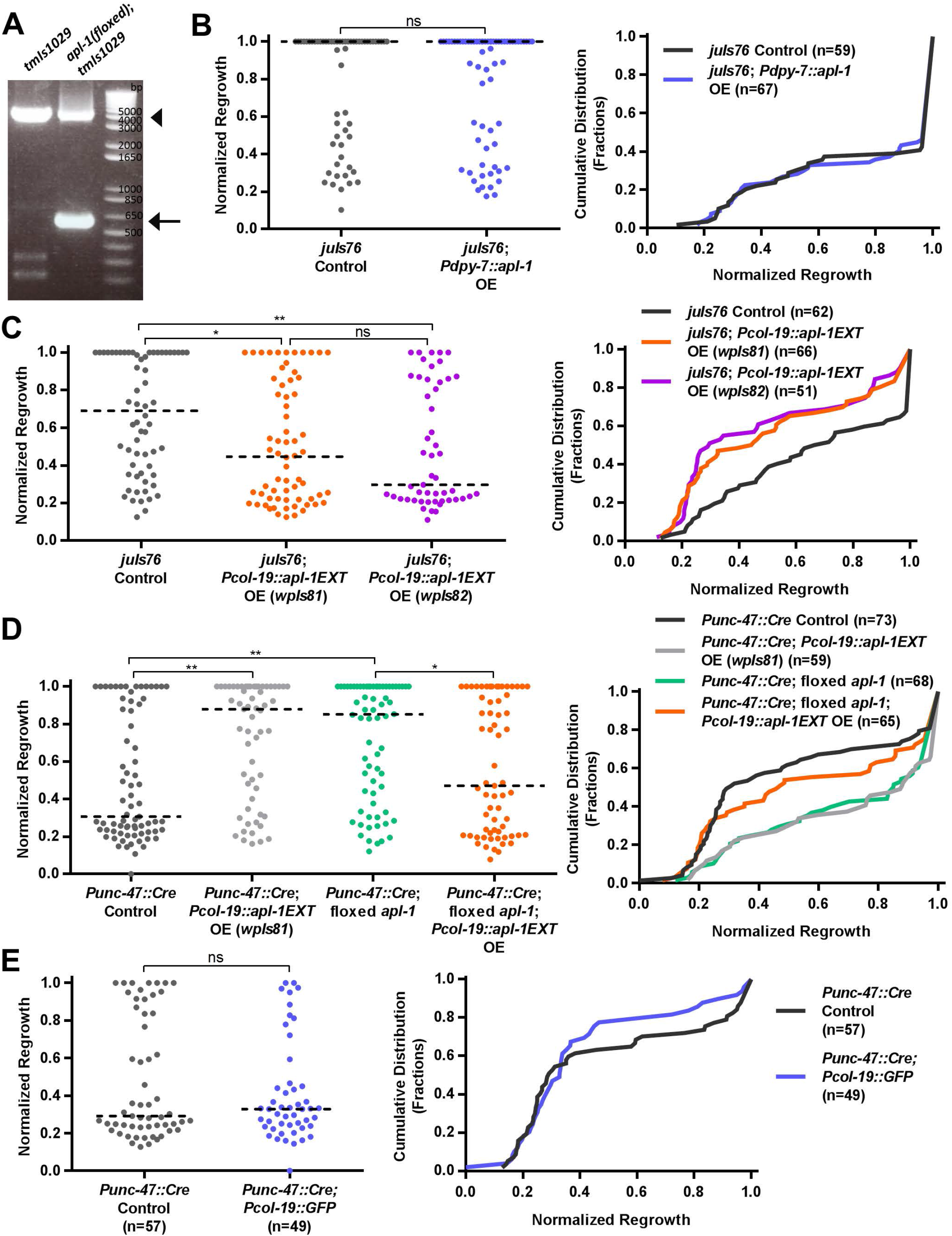
(A) DNA gel of single worm PCR genotyping across the *apl-1* endogenous genomic locus, comparing hypodermal Cre control *(tmls1029)* versus hypodermal specific Cre-loxP apl-1 cKO *[apl-1(wp19);tmls1029].* Arrow head indicates the wildtype size. Arrow indicates the size corresponding to after Cre-loxP excision. (B) Axon regeneration (from L4 to 1 day adult) is not affected by developmental overexpression of *apl-1* from the hypodermis *(dpy-7* promoter, active before L4). (Kolmogorov-Smirnov D=0.05793; P>0.9999) (C) Axon regeneration is decreased in two lines with different integrated arrays both overexpressing secreted APL-1EXT fragment from the hypodermis *(col-19* promoter). *juIs76* control versus *wpIs81* Kolmogorov-Smirnov D=0.2761, P=0.0153. *juIs76* control versus *wpIs82* Kolmogorov-Smirnov D=0.3393, P=0.0032. The effects of the two integrated arrays *(wpIs81* and *wpIs82)* are statistically indistinguishable (Kolmogorov-Smirnov D=0.1221, P=0.7843). (D) APL-1EXT overexpression from the hypodermis *(col-19* promoter, *wpIs81)* inhibits axon regeneration in the *apl-1* GABA-specific cKO background (Kolmogorov-Smirnov D=0.2443; P=0.0378). APL-1EXT overexpression increases regeneration in the Cre only control background which has lower baseline regeneration (Kolmogorov-Smirnov D=0.3316; P=0.0015). *(apl-1* GABA-specific cKO vs Cre only control Kolmogorov-Smirnov D=0.3253, P=0.0012). (E) Axon regeneration is not significantly affected by *Pcol-19*∷*GFP* overexpression in *Punc-47*∷*Cre* control background (Kolmogorov-Smirnov D=0.1583; P=0.5242).

**Figure S6.**
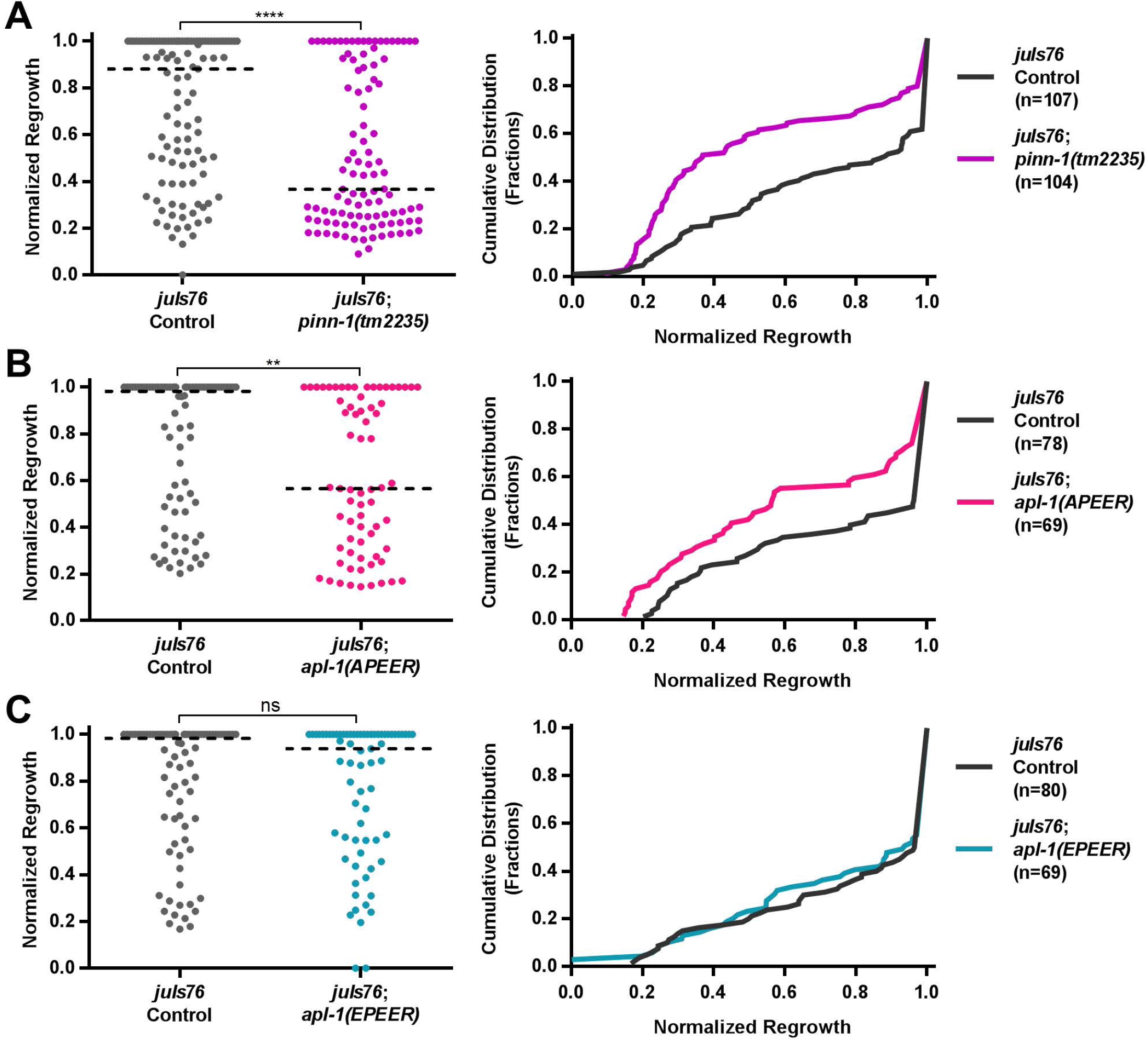
(A) Axon regeneration is decreased in the *pinn-1(tm2235)* mutant compared to the *juIs76* control (Kolmogorov-Smirnov D=0.3158; P<0.0001) (B) Axon regeneration is decreased in the *apl-1(APEER)* mutant compared to the *juIs76* control (Kolmogorov-Smirnov D=0.2776; P=0.0071) (C) Axon regeneration is not affected in the *apl-1(EPEER)* mutant compared to the *juIs76* control (Kolmogorov-Smirnov D=0.08333; P=0.9591

